# Comparative identification of microRNAs in *Apis cerana cerana* workers’ midguts responding to *Nosema ceranae* invasion

**DOI:** 10.1101/528166

**Authors:** Dafu Chen, Yu Du, Huazhi Chen, Haipeng Wang, Cuiling Xiong, Yanzhen Zheng, Chunsheng Hou, Qingyun Diao, Rui Guo

## Abstract

MicroRNAs (miRNAs) are endogenous small noncoding RNAs that post transcriptionally regulate gene expression and are involved in many biological processes including host-pathogen interactions. However, the potential role of miRNAs in the responses of eastern honeybees to *Nosema ceranae* invasion is completely unknown. Here, the expression profiles and differentially expressed miRNAs (DEmiRNAs) in the midguts of *Apis cerana cerana* workers 7 and 10 days post infection (dpi) with *N. ceranae* were investigated via small RNA sequencing and bioinformatics. In total, 529 miRNAs highly conserved between various species and 25 novel miRNAs with varied expressions were identified for the first time. In addition, stem-loop RT-PCR confirmed the expression of 16 predicted miRNAs, validating their existence. Eight up-regulated miRNAs and six down-regulated miRNAs were detected in midguts at 7 dpi, while nine and three miRNAs were significantly up-regulated and down-regulated, respectively, in midguts at 10 dpi. In addition, Venn analysis showed that five DEmiRNAs were shared, while nine and seven DEmiRNAs were specifically expressed in midguts at 7 and 10 dpi, respectively. Gene ontology analysis suggested that a portion of the DEmiRNAs and corresponding target genes were involved in various biological processes, cellular components, and molecular functions including immune system processes and response to stimulus and signaling. Moreover, KEGG pathway analysis shed light on the potential functions of some DEmiRNAs in the regulation of target genes engaged in material and energy metabolism, cellular immunity such as endocytosis and phagosome, and the humoral immune system, including the Jak-STAT and MAPK signaling pathways. Further investigation demonstrated a complex regulation network between DEmiRNAs and their target mRNAs, with miR-598-y, miR-252-y, miR-92-x and miR-3654-y at the center of the network, implying their key parts in host responses. This comprehensive miRNA transcriptome analysis demonstrated that *N. ceranae* invasion influenced the expression of miRNAs in the midguts of *A. c. ceranae* workers; the results can not only facilitate future exploration of the regulatory roles and mechanisms of miRNAs in hosts’ responses, especially their immune responses to *N. ceranae*, but also provide potential candidates for further investigation of the molecular mechanisms underlying eastern honeybee-microsporidian interactions.

## 1. Introduction

Honeybees play vital roles not only in the pollination of crops and wild flora but also in the support of critical ecosystem balance^1–2^. In addition, honeybees serve as key models for studying development, social behavior, and disease transmission^3–4^. European honeybees (*Apis mellifera*) and Asian honeybees (*Apis cerana*), the two best-known honeybee species, have been domesticated for honey production and crop pollination. Compared with *A. mellifera*, *A. cerana* has some prominent advantages, such as long flight duration, adaptation to extreme weather conditions, frequent grooming and hygienic behaviors, and cooperative group-level defenses^5^. *Apis cerana cerana*, a subspecies of *A. cerana*, is an important endemic species that is crucial to ecosystem balance and environmental improvement in China^6^.

Nosemosis is a serious disease in adult honeybees caused by several *Nosema* species. *Nosema* infection occurs through the ingestion of spores in contaminated food or water, followed by the germination of these spores triggered by the physical and chemical conditions of the midgut. The genetic material is then transferred into the host cell of the midgut epithelium where it multiplies, and finally, the spores are excreted from the bee in the feces, which provides new sources of the infection via cleaning and feeding activities in the colonies, or are disseminated into the environment^7–9^. Microsporidia are spore-forming and obligate intracellular fungal pathogens, which can infect a wide variety of hosts, ranging from insects to mammals^10^. *N. ceranae* is a widespread microsporidian pathogen of honeybees, which was first discovered and described from *A. cerana* near Beijing, China^11^, and shortly afterward, it was reported to have infected *A. mellifera* in Europe and Taiwan^12–13^. At present, *N. ceranae* has been identified in colonies of *A. mellifera* all over the world^14–15^. *N. ceranae* is infective to all castes in the colony, including queens, drones and workers^9^. It can remarkably reduce colony strength and productivity but also interacts with other environmental stressors to weaken colony health^16–17^.

MicroRNAs (miRNAs) are single-stranded, highly conserved, noncoding RNA molecules of 19-24 nucleotides in length (nt) that negatively regulate gene expression by targeting specific sites in the 3’ untranslated region (UTR) of mRNAs at the posttranscriptional level or by mediating the degradation of the target gene^18^. Studies over the past years have shown that miRNAs are able to play key parts in diverse biological processes, including cell proliferation^19^, apoptosis^20^, morphogenesis^21^, and metabolism^22^, as well as playing roles in stress^23^ and immune responses^24^.

In comparison to other insects such as *Drosophila* and the silkworm, little research has been done on the roles of miRNAs during the host-parasite interactions in honeybees. Huang and colleagues performed deep sequencing of western honeybee miRNAs daily across the six day reproductive cycle of *N. ceranae* and detected 17 DEmiRNAs that target more than 400 mRNAs involved in several pathways including metabolism^25^. In eastern honeybees infected with *N. ceranae*, a series of changes are accompanied by alterations in gene expression, and miRNAs are likely to be involved in this process. However, there has been no report of studies addressing this issue until now. In the current work, in order to investigate eastern honeybee miRNAs and expand upon novel insights into host-parasite interactions in *A. c. cerana*, we utilized deep sequencing and bioinformatics to analyze the dynamic miRNA expression profiles and differentially expressed miRNAs (DEmiRNAs) of *A. c. cerana* after *N. ceranae* invasion and identified the corresponding target mRNAs and pathways involved. Our results provide novel insights into the host-pathogen interactions between *A. c. cerana* and *N. ceranae* and facilitate further investigation of miRNA function in eastern honeybee responses to *N. ceranae*.

## 2. Materials and methods

### 2. 1. Preparation of *N. ceranae* spores

Foraging bees were collected from a heavily infected colony in an apiary of *A. c. cerana* in Fuzhou city, Fujian province, China. The infected honeybees were kept at −20 °C for 20 min to anesthetize them, followed by purification of fresh spores of *N. ceranae*^26–27^ with some modifications. In brief, the midguts of infected workers were removed and crushed in sterile water and filtered with four layers of gauze to remove tissue debris. Following centrifugation of the filtered suspension at 8000 rpm for 5 min, the supernatant was discarded, and the resuspended pellet was further purified on a discontinuous Percoll gradient (Solarbio, China) containing 5 mL each of 25%, 50%, 75% and 100% Percoll solution. The spore suspension was overlaid onto the gradient and centrifuged at 14000 rpm for 90 min at 4 °C. The spore pellet was carefully extracted with a syringe and then centrifuged again on a discontinuous Percoll gradient to obtain clean spores, which were immediately used for experimental infection of *A. c. cerana* workers. The presence of *N. ceranae* spores and the absence of *Nosema apis* spores were confirmed by PCR using previously described primers^15^.

### 2. 2. Experimental inoculation of honeybees

Frames of a sealed brood from a healthy colony of *A. c. cerana* (*Nosema*-free, as verified by PCR) sited in an experimental apiary of the College of Bee Science, Fujian Agriculture and Forestry University, were swiftly transferred to the laboratory, and kept in an incubator at 34 ± 2 °C to provide newly emerged *Nosema*-free workers. Workers were carefully removed, confined in plastic cages in groups of 30, and kept in an incubator at 34 ± 2 °C. According to the standard method previously described^28^, 24 h after eclosion, the workers were starved for 2 h and then each was fed with 5 μL of a 50% sucrose (w/w in water) solution containing 1×10^5^ *N. ceranae* spores^29^. For *N. ceranae*-treated groups, honeybee workers were held with their mouthparts touching a droplet with the spore solution at the tip of a micropipette until the worker had consumed the entire droplet; workers that did not consume the entire droplet were discarded. For the control group, each worker in three replicates was fed 5 μL of a 50% sucrose solution without *N. ceranae* spores. The honeybees were fed *ad libitum* with a solution of sucrose (50% w/w), and the feeders were replaced daily throughout the whole experiment. Each cage was examined daily, and dead bees were removed and counted. Nine bees from each cage in the *N. ceranae*-treated and control groups were collected at 7 d and 10 d post inoculation (dpi) and then sacrificed, followed by dissection of the intestinal tract and separation of the midgut from the rectum and Malpighian tubules. For each cage, midgut tissues were immediately pooled from all nine workers, frozen in liquid nitrogen and stored at −80 °C until sRNA-seq, stem loop RT-PCR and real time quantitative PCR (RT-qPCR) was performed.

### 2. 3. Small RNA isolation, cDNA library construction and deep sequencing

Firstly, total RNA of each midgut sample in *N. ceranae*-treated and control groups were extracted using TRIzol Reagent (Invitrogen, USA) following the manufacturer's protocols. Secondly, DNA contaminants were removed with RNase-freeDNase I (TaKaRa, China). The purified RNA quantity and quality were checked using Nanodrop 2000 spectrophotometer (Thermo Fisher, USA), and the integrity of the RNA samples were evaluated using Agilent 2100 bioanalyzer (Agilent Technologies, USA) and only values of 28S/18S≥0.7 and RIN≥7.0 were considered qualified for the subsequent small RNA library construction. Thirdly, RNA molecules in the size range of 18-30 nt were enriched by agarose gel electrophoresis and then ligated with 3’ and 5’ RNA adaptors, and fragments with adaptors on both ends were enriched by PCR after reverse transcription. Fourthly, the subsequent cDNAs were purified and enriched by 3.5% AGE to isolate the expected size (140-160 bp) fractions and eliminate unincorporated primers, primer dimer products, and dimerized adaptors. Ultimately, the 12 cDNA libraries were sequenced on Illumina sequencing platform (HiSeq™ 4000) using the single-end technology by GENE DENOVO Biotechnology Co. (Guangzhou, China).

### 2. 4. Quality control and sequencing data analysis

The raw data were pre-processed to exclude low-quality reads (length <20 nt and ambiguous N), 5′ adapter, 3′ adapter and poly(A) sequences to gain clean reads, which were aligned against NCBI GeneBank and Rfam databases to remove noncoding RNA including rRNA, scRNA, snoRNA, snRNA and tRNA. Then, the obtained sequences were compared with exons and introns in the *A. cerana* genome (assembly ACSNU-2.0) to classify mRNA degradation products and the repeat associate miRNA sequences. All the downstream analyses were based on clean reads with high quality.

By utilizing Bowtie (v 1.1.0)^30^, the filtered sequences were analyzed with BLAST search against miRBase 21.0 by allowing at most two mismatches outside of the seed region^31^, and small RNAs that matched known miRNAs of other animal species in miRBase were identified as conserved (known) miRNAs. The sequences that did not match conserved miRNAs were used to identify potentially novel miRNA candidates using RNAfold software^32^. Only sequences with typical stem-loop hairpins and free energy lower than −20 kcal/mol were considered as potential novel miRNAs. Size distribution and saturation analysis were performed using sequencing libraries after all annotation steps. Hierarchical clustering of novel miRNA expression was conducted using heatmap tool in OmicShare (http://www.omicshare.com/tools/).

### 2. 5. Analysis of DEmiRNAs

The miRNA expression levels in each sample were normalized to the total number of sequence tags per million (TPM) following formula: normalized expression = mapped read count/total reads × 10^6^. Differential expression analysis of two samples was performed using the DEGseq R package^33^ and *p* values were adjusted using *q* value. The criteria of *p* value (FDR) < 0.05 and |log_2_(Fold change)| > 1 were set as the threshold for statistically significant differential expression. A positive value indicated up-regulation of a miRNA, while a negative value indicated down-regulation.

### 2. 6. Prediction of the target genes of DEmiRNAs and construction of miRNA-mRNA regulation networks

Target genes for miRNAs were predicted with miRanda (v3.3a)^34^, RNAhybrid (v2.1.2)+svm_light (v6.01)^35^ and TargetFinder (Version: 7.0)^36^ software. The input files were miRNA FASTA sequences files. Intersections of the results from the three abovementioned programs comprised the final predicted gene targets. Then, miRNA-mRNA regulation networks were visualized using Cytoscape v.3.2.1 software^37^.

### 2. 7. GO and KEGG pathway analyses of DEmiRNA target genes

To determine their main biological functions, DEmiRNAs target genes were annotated by terms in the Gene Ontology (GO) database (http://www.geneontology.org/) using Blast2GO^38^, following their numeric orders in the NCBI nr database. Subsequently, GO functional categorizations for all target genes were determined with WEGO software^39^, and GO enrichment analysis of functional significance terms in the GO database was performed using the hypergeometric test to identify significantly enriched GO terms for the target genes compared to the genome background. KEGG pathway analyses of the predicted target genes were performed using the KEGG pathway database (http://www.genome.jp/kegg/pathway.html)^40^.

### 2. 8. Stem-loop RT-PCR confirmation of miRNAs

Total RNAs from AcCK1 and AcCK2 samples were isolated using RNAiso plus kit (TaKaRa, China) and then treated with DNase I (TaKaRa, China) to remove remaining DNA. According to the previously described method^41^, stem-loop primers, specific forward primers and universal reverse primers (presented in **Table S1**) were designed using DNAMAN software on basis of the sequences of the randomly selected nine *A. c. cerana* miRNAs, including miR-1943-x, miR-3793-x, miR-252-y, miR-3963-x, miR-8516-x, miR-149-y, miR-7311-y, miR-9008-x, novel-m0014-3p. The primers were synthesized by Sangon Biotech Co., Ltd (Shanghai, China). One microgram of total RNA was reverse transcribed to cDNA using RevertAid First Strand cDNA Synthesis Kit (TaKaRa, China) and stem-loop primers (**Table S1**). The PCR amplification of randomly selected miRNAs was conducted on a T100 thermo cycler (BIO-RAD) using Premix (TaKaRa, China) under the following conditions: pre-denaturation step at 94 °C for 5 min; 30 amplification cycles of denaturation at 94 °C for 50 s, annealing at 55 °C for 30 s, and elongation at 72 °C for 1 min; followed by a final elongation step at 72 °C for 10 min. The PCR products were detected on 2% agarose gel eletrophoresis with Genecolor (Gene-Bio, China) staining.

### 2. 9. RT-qPCR validation of DEmiRNAs

DEmiRNAs including miR-6547-x, miR-7311-x, miR-2779-y, miR-3726-x, miR-1788-y, miR-3319-y, miR-1672-x, novel-m0005-3p, and novel-m0008-3p were randomly selected for RT-qPCR validation. Stem-loop primers and specific forward primers (presented in **Table S1**) were designed and synthesized using the aforementioned method. cDNA Synthesis was conducted using stem-loop primer following the method mentioned above. RT-qPCR was carried out in an Applied Biosystems QuantStudio 3 (Thermo Fisher, China) in a 20 μl reaction volume containing 1 μl cDNA, 10 μl SYBR Premix (Vazyme, China), 0.2 μl specific forward primer (20 μM), 0.2 μl reverse primer (20 μM), and 8.6 μl DEPC water. The reaction was performed at 95°C for 5 min, followed by 45 cycles of 94°C for 15 s, 60°C for 15 s, and 72°C for 15 s. The abundance of miRNAs was normalized relative to that of endogenous control snRNA U6. All reactions were performed in triplicate. The threshold cycle (Ct) was determined using the default threshold settings, and the data were analyzed using 2^−ΔΔCt^ program^42^. The experiment was performed independently three times.

## 3. Results

### 3. 1. Overview of the small RNA library from *A. c. ceranae* workers’ guts

To identify the miRNA profiles and DEmiRNAs during the responses of an *A. c. ceranae* worker’s midgut to *N. ceranae* invasion, 12 small RNA libraries were constructed using Illumina sequencing. In total, 127,523,419 raw reads with an average of ~106,269,512 reads per sample were produced, and after filtering out low-quality sequences, 5′ and 3′ adaptors and reads 18<nt, 122,104,443 clean reads were obtained for further analysis (**Table 1**). In addition, the Pearson correlation between every sample in each group was above 0.9619 (**Figure S1**), suggesting sufficient reproducibility and rationality of sampling. A BLAST run against the NCBI GenBank and RFam databases identified 3,611,375 (36.20%) to 5,054,110 (52.00%) unique small RNAs as rRNA, 101,602 (1.02%) to 221,407 (1.97%) as tRNAs, 5,123 (0.05%) to 11,137 (0.11%) as snRNAs, and 176 (0.002%) to 584 (0.005%) as snoRNAs (**Table 2**).

**Table 1.** Summary of small RNA sequencing datasets from the *N. ceranae*-treated and untreated midgut samples of *A. c. cerana*.

| Samples | Raw reads | Clean reads | Clean tags |
| --- | --- | --- | --- |
| AcCK1-1 | 12757706 | 12049987 (94.4526%) | 10213625 (84.7605%) |
| AcCK1-2 | 12794543 | 12122624 (94.7484%) | 10275851 (84.7659%) |
| AcCK1-3 | 11328883 | 10727931 (94.6954%) | 8671328 (80.8295%) |
| AcCK2-1 | 17161911 | 16292122 (94.9319%) | 14528074 (89.1724%) |
| AcCK2-2 | 11666305 | 11306117 (96.9126%) | 9862922 (87.2353%) |
| AcCK2-3 | 11757223 | 11328016 (96.3494%) | 9310382 (82.1890%) |
| AcT1-1 | 11213906 | 10923950 (97.4143%) | 9356024 (85.6469%) |
| AcT1-2 | 14549245 | 13778004 (94.6991%) | 11661491 (84.6385%) |
| AcT1-3 | 11029767 | 10800311 (97.9197%) | 8914299 (82.5374%) |
| AcT2-1 | 13263930 | 12775381 (96.3167%) | 10974972 (85.9072%) |
| AcT2-2 | 13688082 | 13150156 (96.0701%) | 10977135 (83.4753%) |
| AcT2-3 | 12936357 | 12467737 (96.3775%) | 10859224 (87.0986%) |

**Table 2.** Small RNA profiling and distribution among the four *A. c. cerana* workers’ midgut samples.

| Types | AmCK1 | AmT1 | AmCK2 | AmT2 |
| --- | --- | --- | --- | --- |
| Total RNA | 9720268 | 9977271 | 11233793 | 10937110 |
| miRNA | 978435 | 577121 | 608488 | 286809 |
| tRNA | 192134 | 101602 | 221407 | 146370 |
| rRNA | 5054110 | 3611375 | 4923233 | 4631227 |
| snRNA | 11137 | 5901 | 6620 | 5123 |
| snoRNA | 244 | 332 | 176 | 584 |
| Exon:+ | 15918 | 6798 | 9246 | 11943 |
| Exon:- | 88024 | 51035 | 40587 | 26733 |
| Intron:+ | 19686 | 10076 | 14761 | 12624 |
| Intron:- | 2979 | 2330 | 2681 | 3447 |
| Repeat | 457936 | 293249 | 415936 | 415268 |
| Unannotated | 3300268 | 5574461 | 5370922 | 5793014 |

Base composition, which is a fundamental feature of miRNA sequences, influences miRNA physiochemical and biochemical properties^43–44^. The sequence lengths of the clean reads in the 12 small RNA libraries were almost all distributed between 16-30 nt, with peaks at 22 nt accounting for 14.02% of all clean reads, followed by 20 nt and 21 nt, which accounted for 13.50% and 13.68% of clean reads, respectively (**Figure 1A**). The nucleotide bias of miRNAs in the midgut of the *A. c. ceranae* worker was further investigated, and the results showed that U was the most common nucleotide at the 5’ end of miRNAs (**Figure 1B**), which was consistent with other studies that also found U to be the most common base at the extreme 5’ end of miRNAs^45–47^. Additionally, analysis of the nucleotide bias at each position showed that U and A were mainly located at the beginning and end of reads of *A. c. ceranae* miRNAs (**Figure 1C**), suggesting that AU base pairing may affect miRNA secondary structure or target recognition^48^.

**Figure 1.**
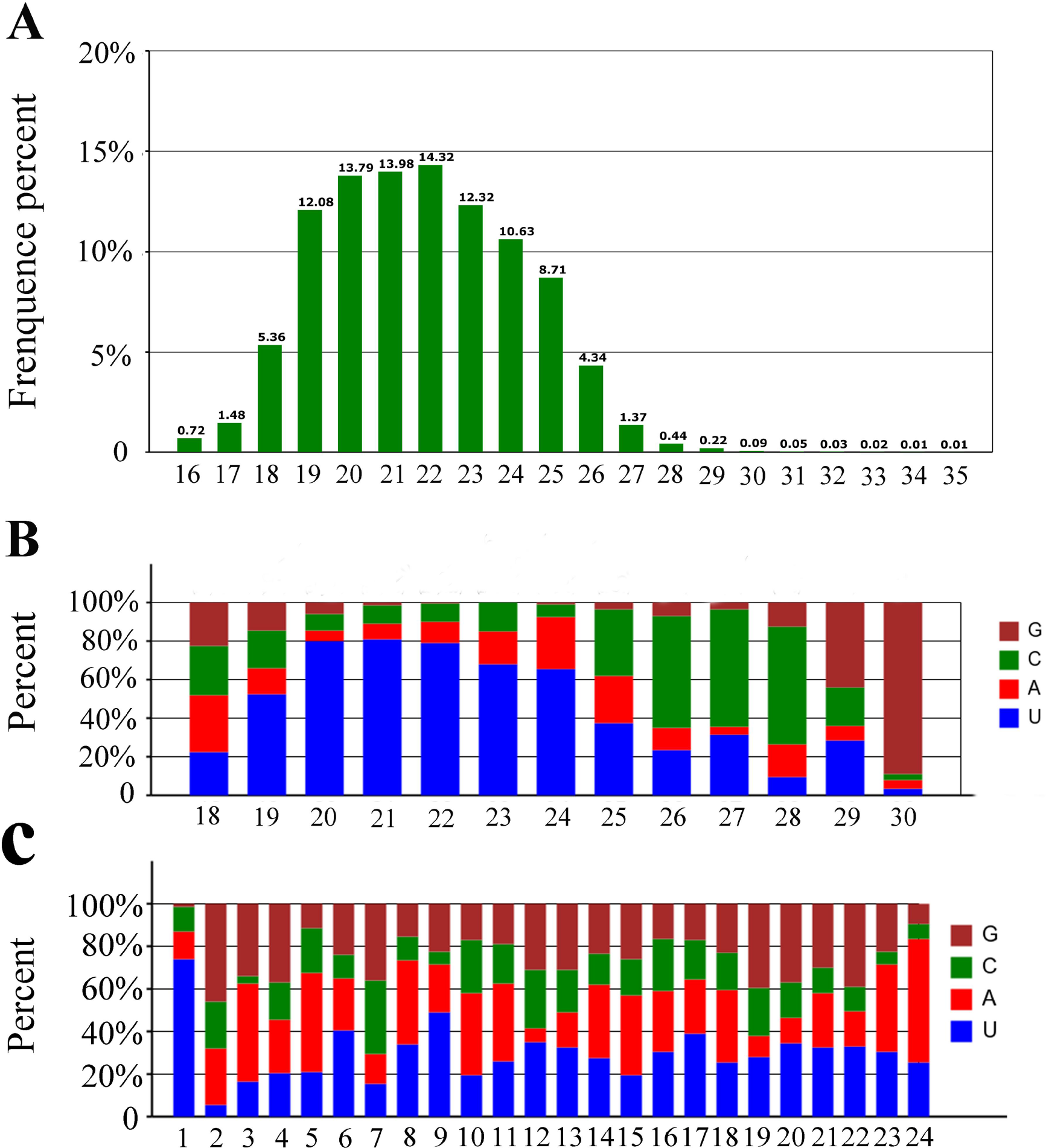
Length distribution and nucleotide bias of the total miRNAs in the untreated and *N. ceranae*-treated *A. c. cerana* workers’ midguts. (A) Length distribution; (B) First nucleotide bias; (C) Nucleotide bias at each position.

### 3. 2. Identification of known and novel miRNAs in *A. c. ceranae* worker’s gut

To explore known miRNAs and novel miRNAs in the four groups (AmCK1, AmT1, AmCK2, AmT2), the mapped sequences were compared to reference genomes and aligned with known mature miRNAs in the miRBase 21.0 database. In total, 340, 338, 306, and 302 conserved miRNAs were identified from the AmCK1, AmT1, AmCK2 and AmT2 samples, respectively, which were highly conserved in other species. These known miRNAs had a broad range of expression levels in *A. c. cerana*, ranging from TPM 90513.14 to TPM 0.3773. Among them, bantam-y, miR-184-y, miR-1-y, miR-276-y and miR-750-y were the most abundant known miRNAs in both AcCK1 and AcCK2 (**Table 3**). We also predicted 25 novel miRNAs not previously found in *A. c. cerana*, including 25, 22, 23 and 17 in AmCK1, AmT1, AmCK2 and AmT2 (**Table 4**), respectively. Among these, novel-m0019-3p, novel-m0001-3p, novel-m0012-5p, novel-m0004-3p and novel-m0012-3p were the highest-expressed novel miRNAs (**Table 4**). Moreover, expression clustering analysis showed that these novel miRNAs had various expression levels in different groups (**Figure 2**). Interestingly, 242 known miRNAs and 23 novel miRNAs were shared by AmCK1 and AmCK2, implying their developmental stage-specific functions. Secondary structures of precursors of three known miRNAs (miR-7-x, miR-9895-y and miR-750-y) and three novel miRNAs (novel-m0001-3p, novel-m0004-3p and novel-m0019-5p) are displayed in **Figure 3**, and they all have classical stem-loop structures.

**Table 3.**
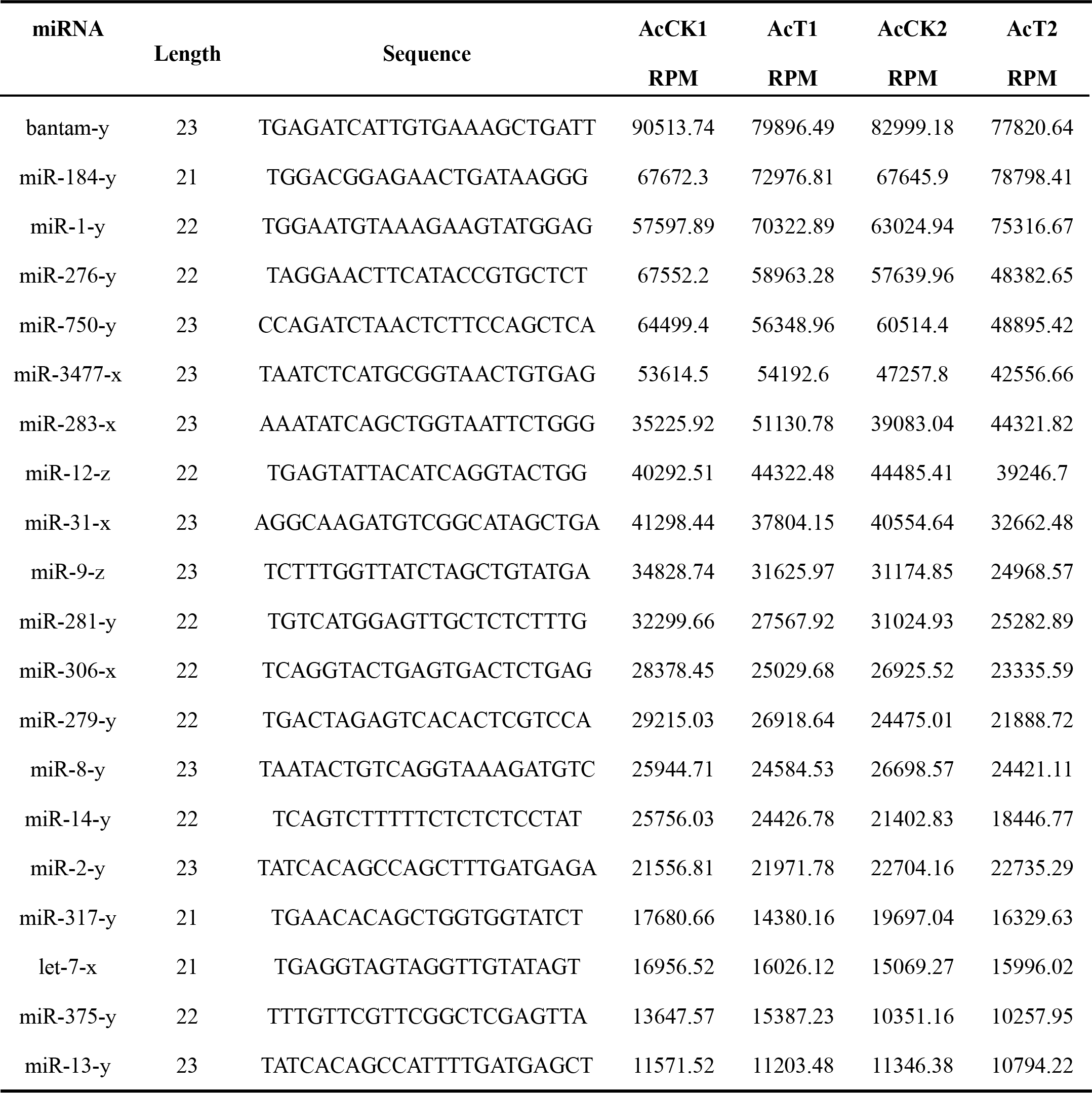
Top 20 highest-expressed known miRNAs in *A. c. cerana* workers’ midguts.

**Table 4.**
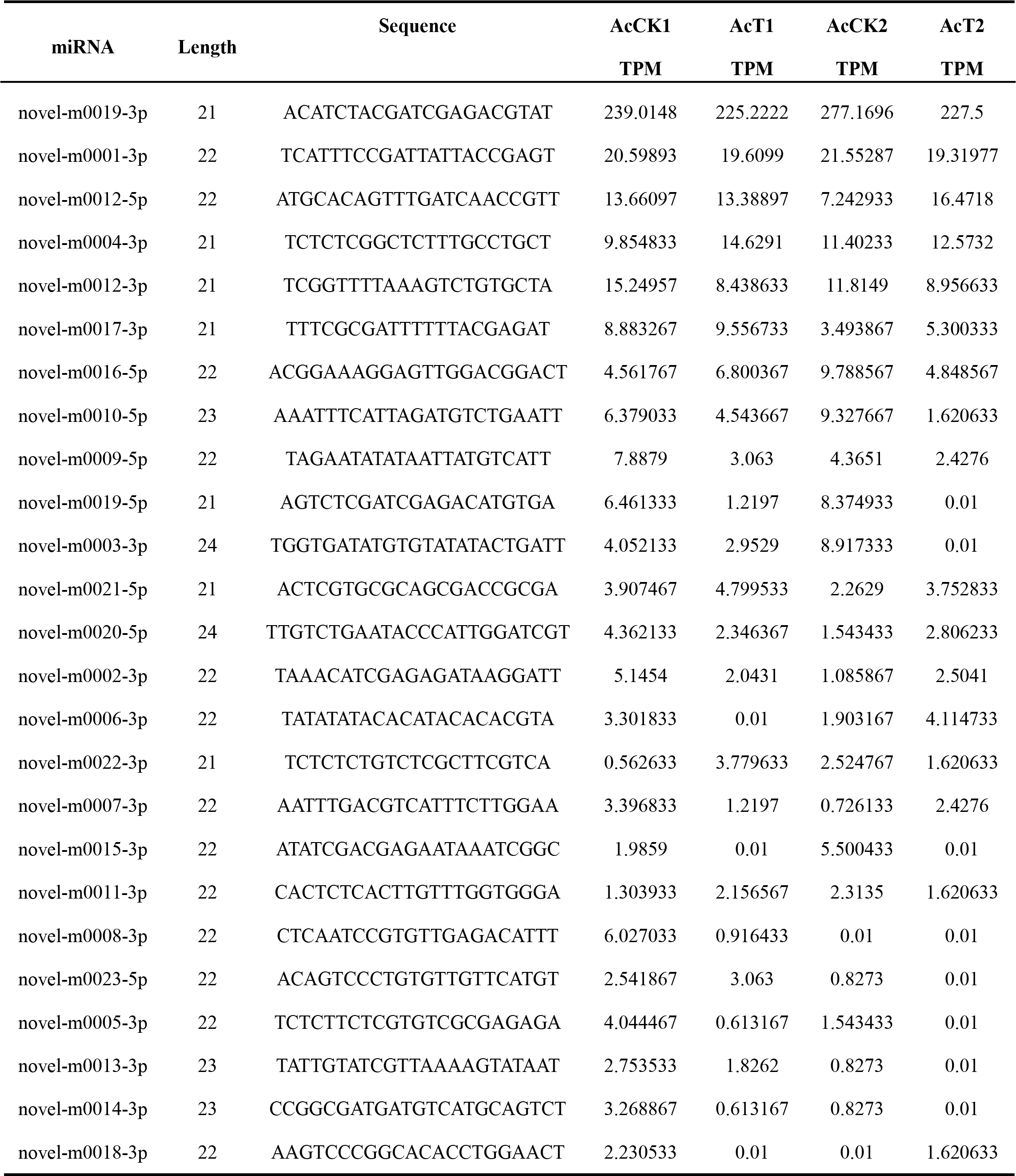
The expressed novel miRNAs in *A. c. cerana* workers’ midguts.

**Figure 2.**
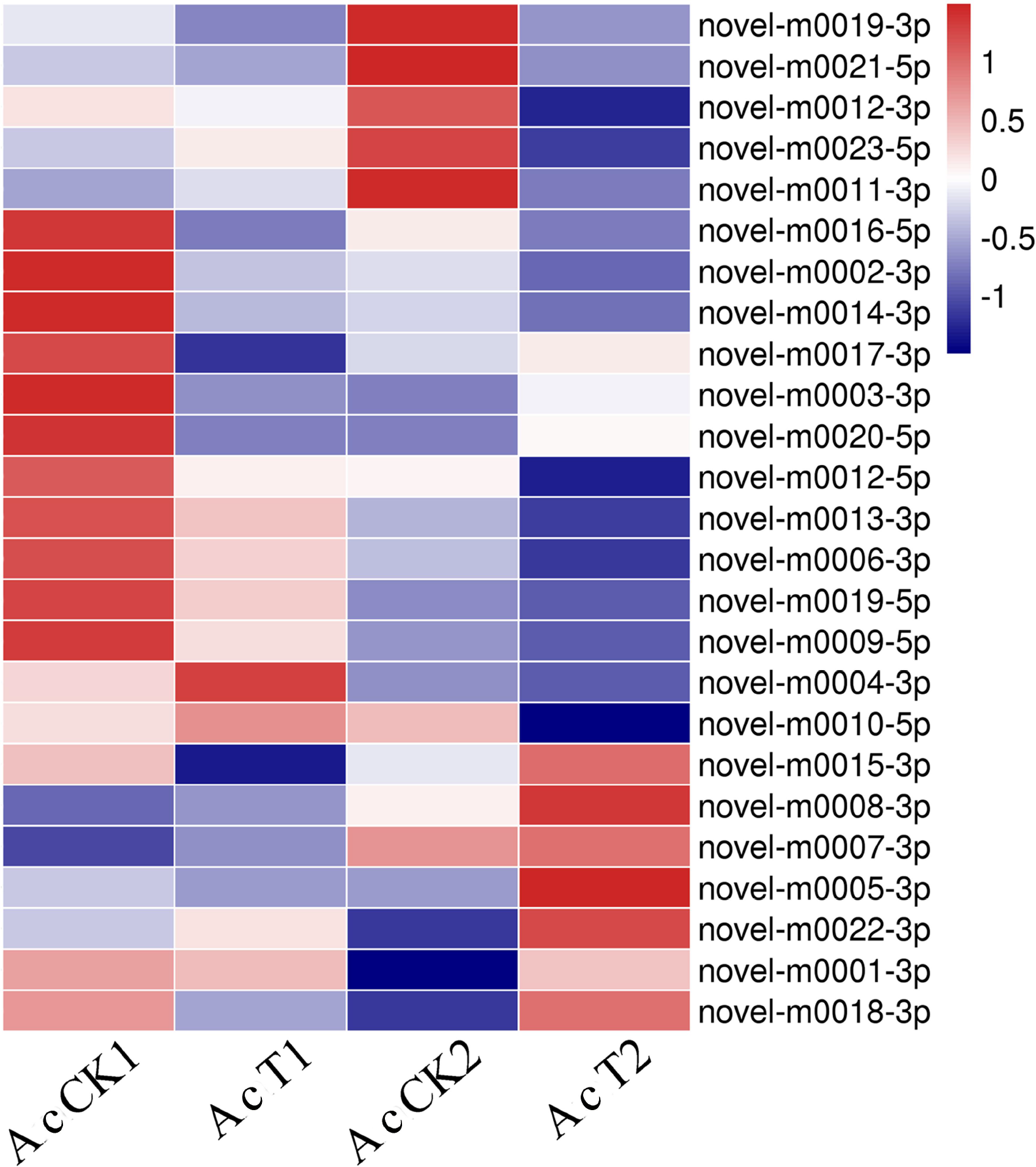
Expression clustering of novel miRNAs in the untreated and *N. ceranae*-treated midguts of *A. c. cerana* workers.

**Figure 3.**
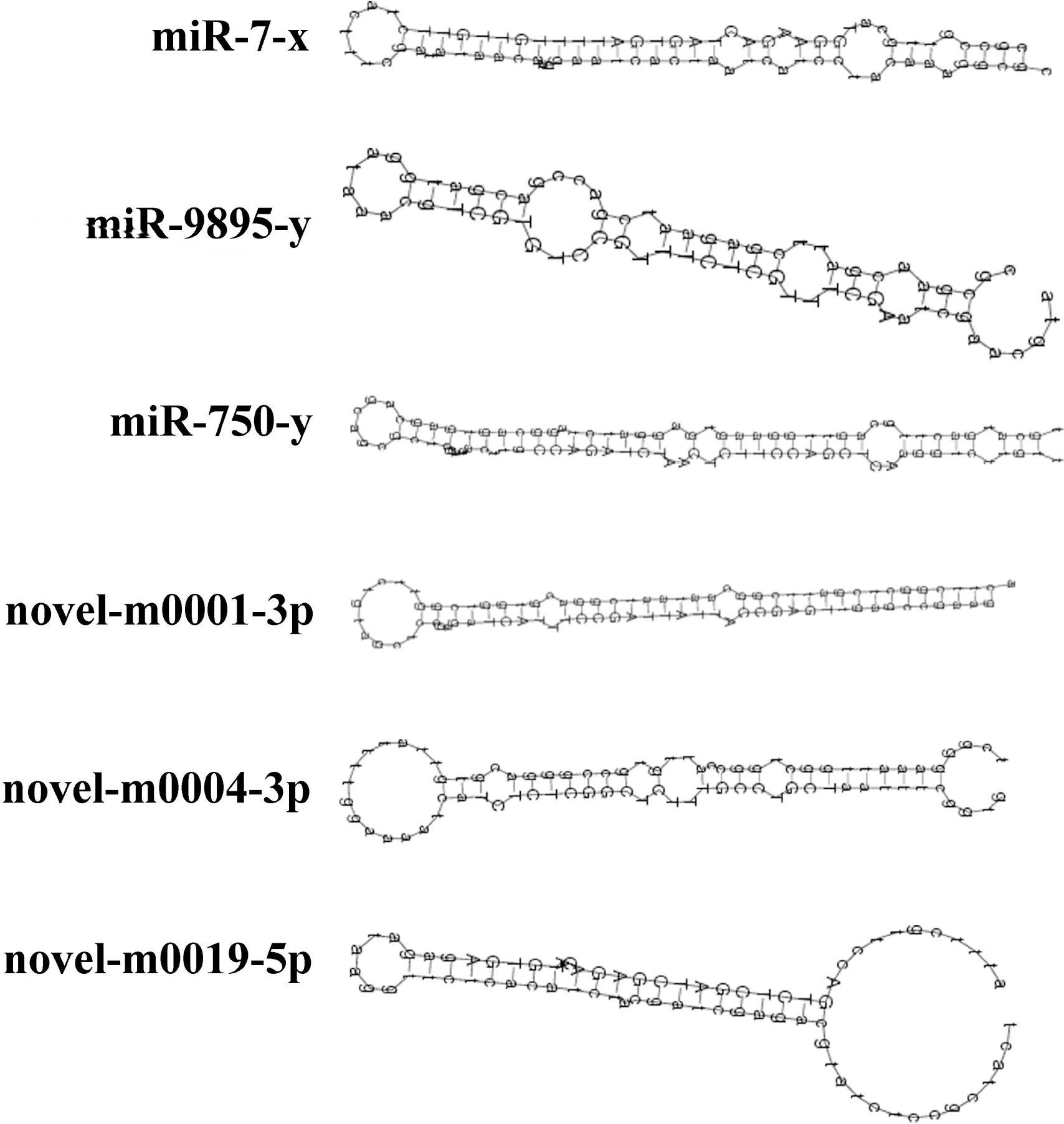
Secondary structure of *A. c. cerana* miRNA precursor. Stem-loop structures of the precursors of miR-7-x, miR-9895-y, miR-750-y, novel-m0001-3p, novel-m0004-3p, and novel-m0009-5p are shown.

Furthermore, nine miRNAs were selected for stem-loop RT-PCR validation, and the results showed that eight of them yielded signal bands in both the AcCK1 and AcCK2 samples (**Figure 4**), indicating that most predicted miRNAs were expressed in the *A. c. cerana* worker’s midgut.

**Figure 4.**
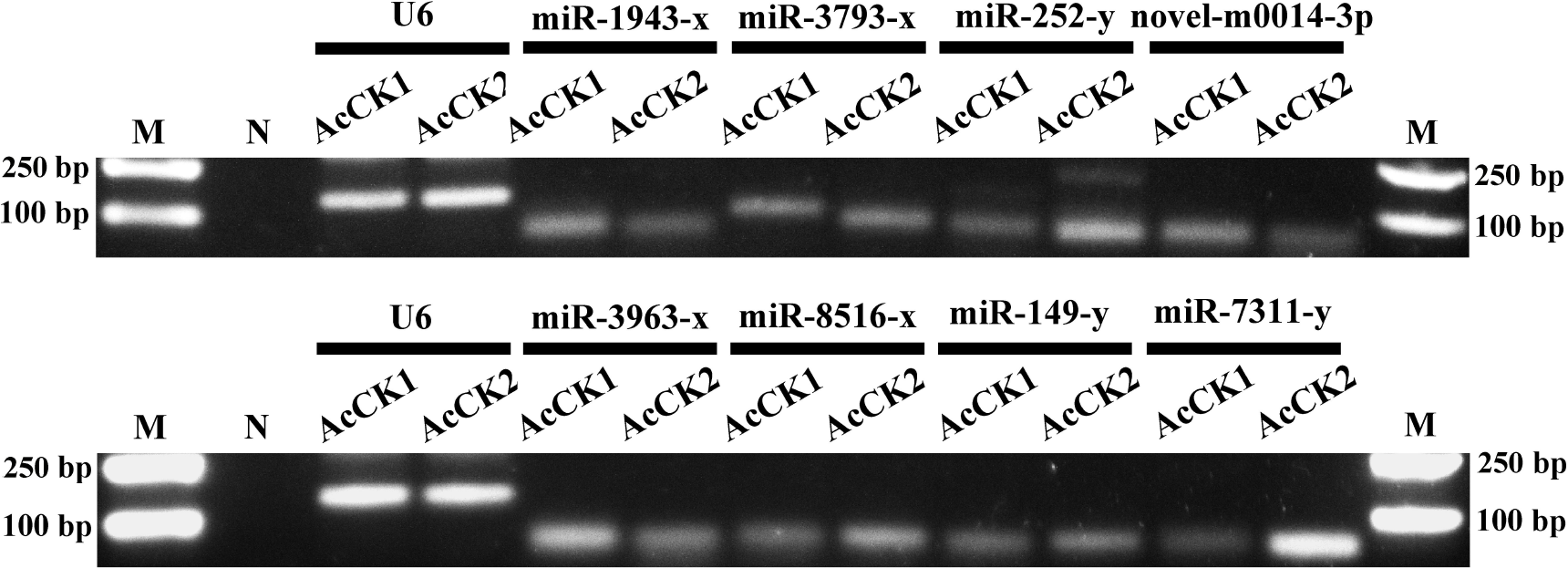
Stem-loop RT-PCR confirmation of eight *A. c. cerana* miRNAs. Lane M: DNA marker; Lane N: Negative control (sterile water was used as PCR template); snRNA U6 was used as positive control.

### 3. 3. Differentially expressed *A. c. ceranae* miRNAs induced by *N. ceranae* invasion

A total of 14 DEmiRNAs were detected in AcT1 VS AcCK1 (**Table S2**), including 8 up-regulated and 6 down-regulated miRNAs, while 12 miRNA with differential expression were identified in AcT2 VS AcCK2 (**Table S3**), including 9 up-regulated and 3 down-regulated ones (**Figure 5A**). Venn diagram analysis was performed, and the result suggested that five DEmiRNAs, including miR-60-y and miR-8462-x, were shared in AcT1 VS AcCK1 and AcT2 VS AcCK2, while nine (eg., miR-1-x and miR-980-y) and seven (eg., novel-m0003-3p and miR-92-x) DEmiRNAs were unique for the two comparison groups, respectively (**Figure 5B**, see also **Table S2** and **Table S3**). In addition, more than 85.71% (12) of the DEmiRNAs in AcT1 VS AcCK1 showed a >7-fold difference in gene expression (**Table S2**), whereas approximately 91.67% (11) of the DEmiRNAs in AcT2 VS AcCK2 displayed a >9-fold differential expression (**Table S3**). We speculate that these DEmiRNAs are likely to play key roles in the regulation of host responses to *N. ceranae* infection.

**Figure 5.**
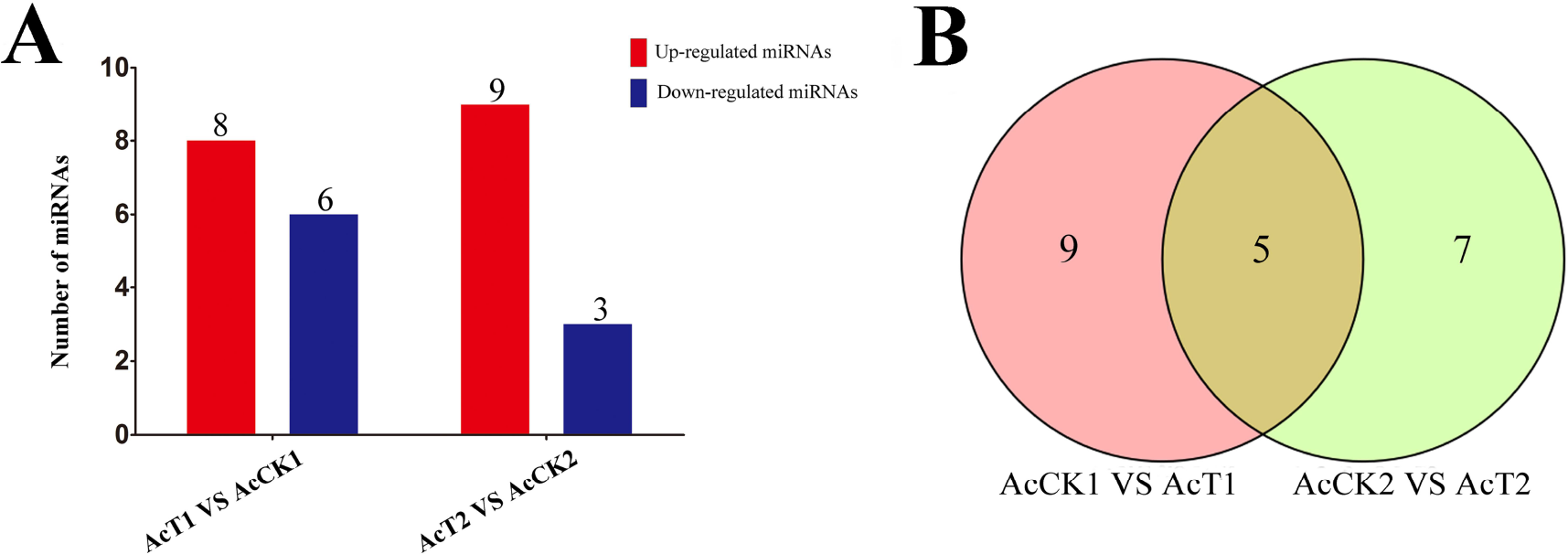
Analysis of DEmiRNAs in *A. c. cerana* workers’ midguts invaded by *N. ceranae*. (A) Summary of the number of DEmiRNAs in AcT1 VS AcCK1 and AcT2 VS AcCK2; (B) Venn diagram of DEmiRNAs in AcT1 VS AcCK1 and AcT2 VS AcCK2.

### 3. 4. Prediction and enrichment analysis of the target genes of *A. c. ceranae* DEmiRNAs

DEmiRNAs in AcT1 VS AcCK1 and AcT2 VS AcCK2 were respectively used to search *A. c. ceranae* 3′-UTR sequences for predicting potential target genes via a combination of RNAhybrid (v2.1.2)+svm_light (v6.01), Miranda (v3.3a), and TargetScan (Version: 7.0) softwares. In total, 2615 target genes of DEmiRNAs were predicted from AcT1 VS AcCK1, while 1905 DEmiRNA target genes in AcT2 VS AcCK2 were identified. GO classification was conducted, and the results indicated that the target genes of DEmiRNAs in AcT1 VS AcCK1 and AcT2 VS AcCK2 were involved in various biological processes (16 and 15 terms), cellular components (11 and 11 terms), and molecular functions (7 and 7 terms) (**Figure 6A, B**, see also **Table S4** and **Table S5**). Cellular process and single-organism process were the most enriched classes in the biological processes; cell and cell part were highly represented in the cellular component categories, and binding and catalytic activity were the top enriched items in molecular functions (**Table S4** and **Table S5**). Further investigation indicated that a large quantity of the shared DEmiRNA target genes were engaged in binding, cellular process, single-organism process, metabolic process, biological regulation, and catalytic activity (**Table S4** and **Table S5**). However, target genes with the function of extracellular matrix, multiorganism process and immune system process were related only in AcT1 VS AcCK1, accounting for three DEmiRNAs including miR-1-x, miR-965-x, and miR-252-y. Supramolecular fiber function was associated only with AcT2 VS AcCK2, accounting for novel-m0003-3p.

**Figure 6.**
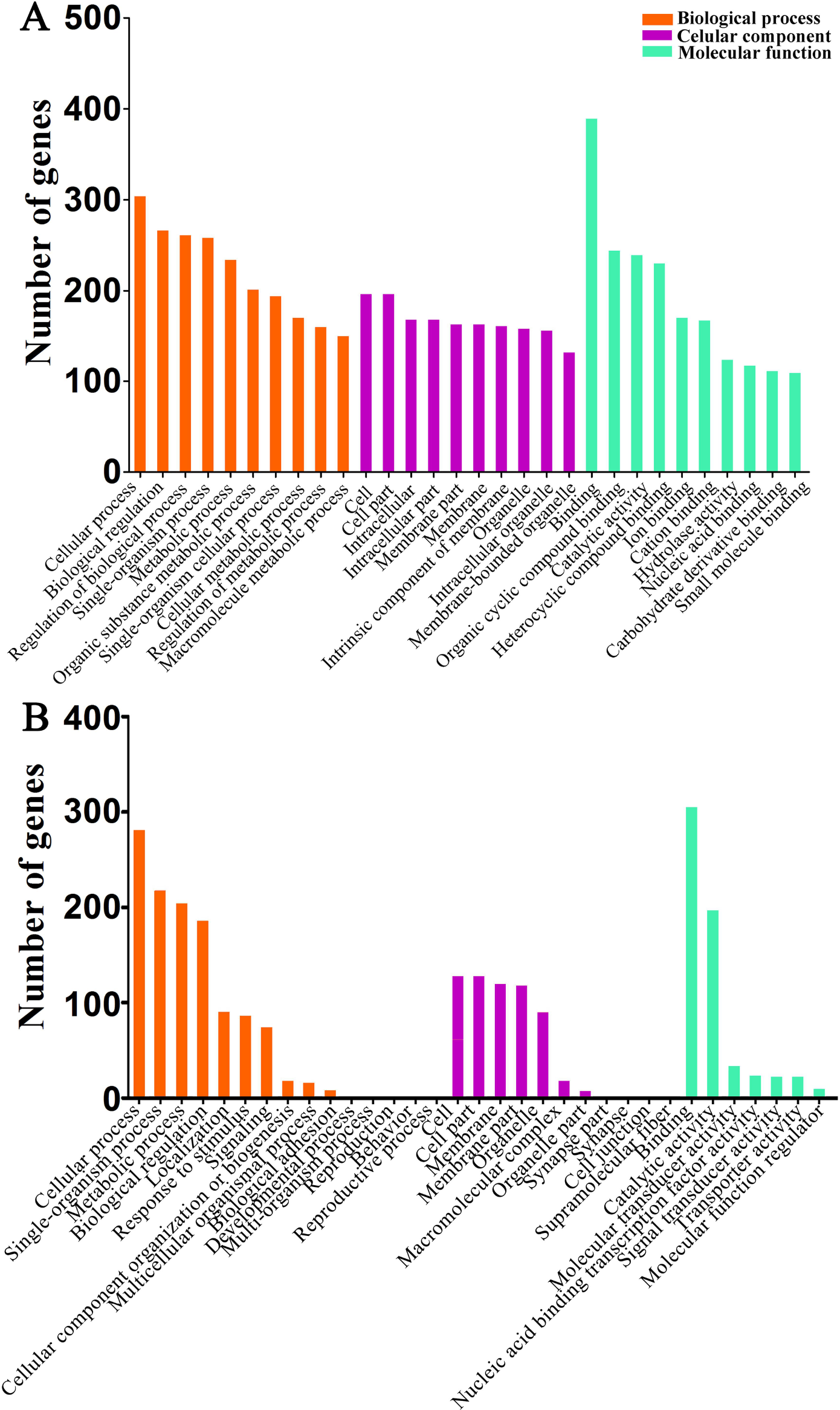
GO categorization of *A. c. cerana* DEmiRNA target genes. (A) Target genes in AcT1 VS AcCK1; (B) Target genes in AcT2 VS AcCK2.

KEGG enrichment analysis for DEmiRNA-targeted genes was conducted, and the results demonstrated that the target genes of shared DEmiRNAs in AcT1 VS AcCK1 were connected with 104 pathways; among them, the highest enriched pathways were endocytosis, the phosphatidylinositol signaling system, the Wnt signaling pathway, inositol phosphate metabolism, and neuroactive ligand-receptor interaction (**Figure 7A, Table S6**). The target genes of DEmiRNAs in AcT2 VS AcCK2 were involved in 92 pathways, and the most enriched pathway was the phosphatidylinositol signaling system, followed by the Wnt signaling pathway, phototransduction, endocytosis and the FoxO signaling pathway (**Figure 7B, Table S7**). Interestingly, 62 and 52 metabolism-related pathways were respectively enriched by 234 and 195 target genes in AcT1 VS AcCK1 and AcT2 VS AcCK2, and, in addition, there were five and six target genes enriched in development in these two comparison groups (**Table S6** and **Table S7**). These results demonstrated that the growth, development and metabolism of *A. c. cerana* workers’ midguts were influenced by corresponding DEmiRNAs via negative regulation.

**Figure 7.**
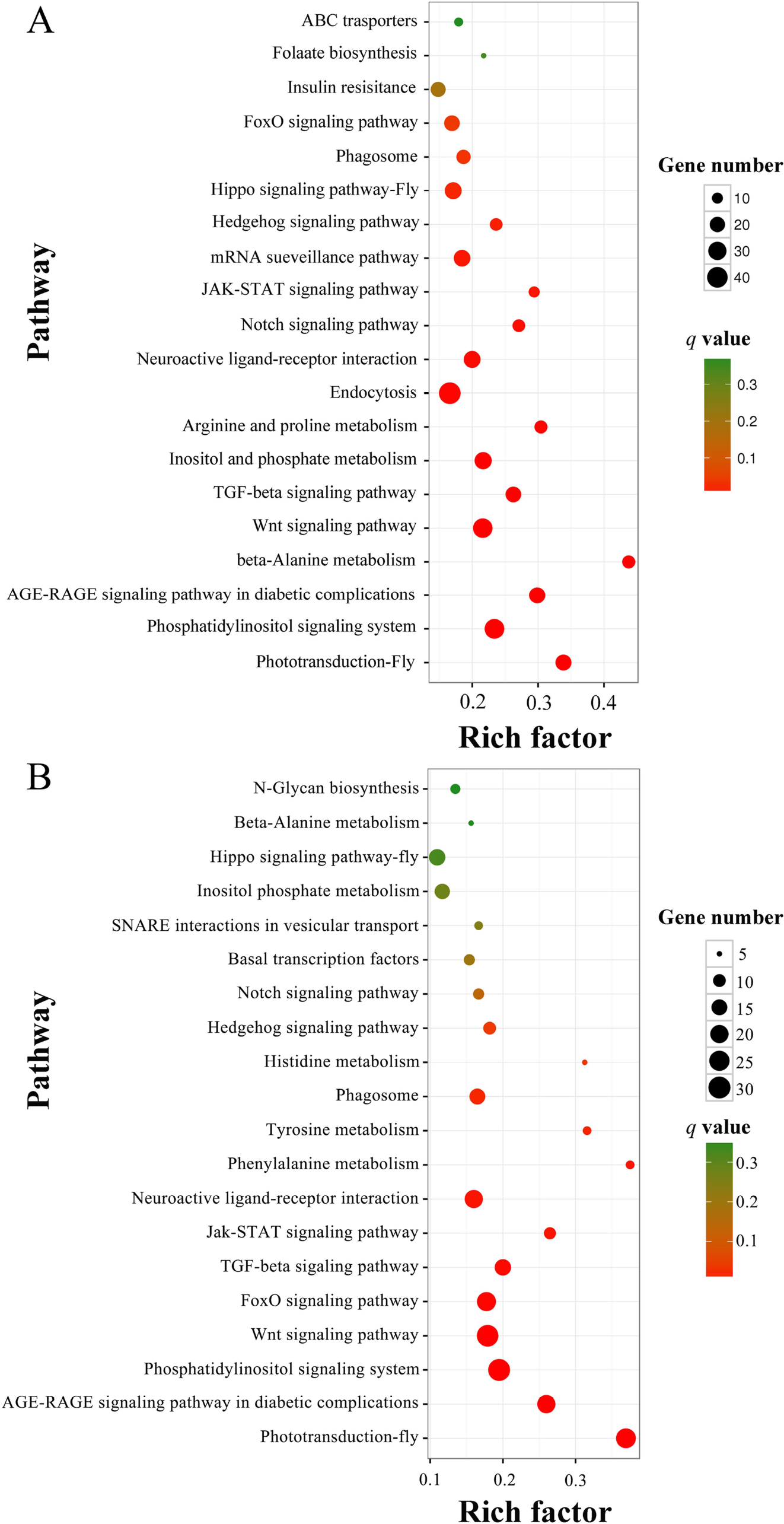
KEGG pathway enrichment analysis for *A. c. cerana* DEmiRNA target genes. (A) Target genes in AcT1 VS AcCK1; (B) Target genes in AcT2 VS AcCK2.

### 3. 5. Regulation networks between *A. c. ceranae* DEmiRNAs and their target genes

Cytoscape was employed for visualization of the regulation networks between DEmiRNA and their target mRNAs. A single mRNA can be targeted by various miRNAs, whereas a single miRNA is also capable of targeting different mRNAs^49^. In these regulation networks, it can be clearly seen that DEmiRNAs and their target genes in AcT1 VS AcCK1 formed complex networks. miR-598-y had as many as 45 target genes, and miR-252-y bound to three targets including XM_017049297.1, XM_017049298.1 and XM_017058024.1 (**Figure 8A**). In contrast, miR-6313-y, miR-3726-x, miR-9204-x and miR-6717-x could bind to only one target (**Figure 8A**). In addition, as shown in **Figure 8B**, for DEmiRNAs in AcT2 VS AcCK2, miR-92-x, miR-3654-y, novel-m0003-3p and miR-6313-y linked to six, four, two and one target genes, respectively.

**Figure 8.**
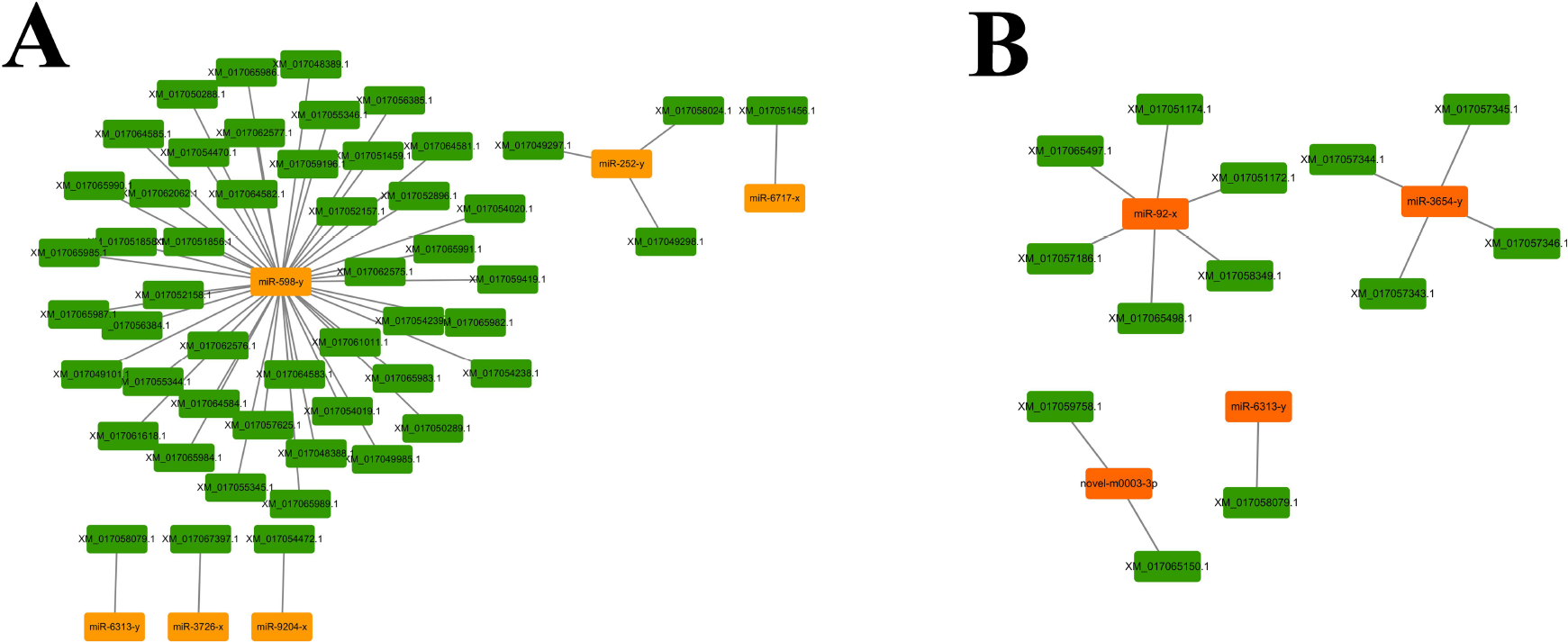
Regulation networks between *A. c. cerana* DEmiRNA and their target genes. (A) DEmiRNA-target gene regulation network in AcT1 VS AcCK1; (B) DEmiRNA-target gene regulation network in AcT2 VS AcCK2.

### 3. 6. Verification of *A. c. cerana* DEmiRNAs via RT-qPCR

To verify the high-throughput sequencing data in our study, the expression levels of nine randomly selected DEmiRNAs in AcT1 VS AcCK1 and AcT2 VS AcCK2 were examined by RT-qPCR. The expression trends of eight DEmiRNAs in the two comparison groups were roughly the same as the result of the sRNA-seq (**Figure 9**), indicative of the accuracy and reliability of our sequencing data.

**Figure 9.**
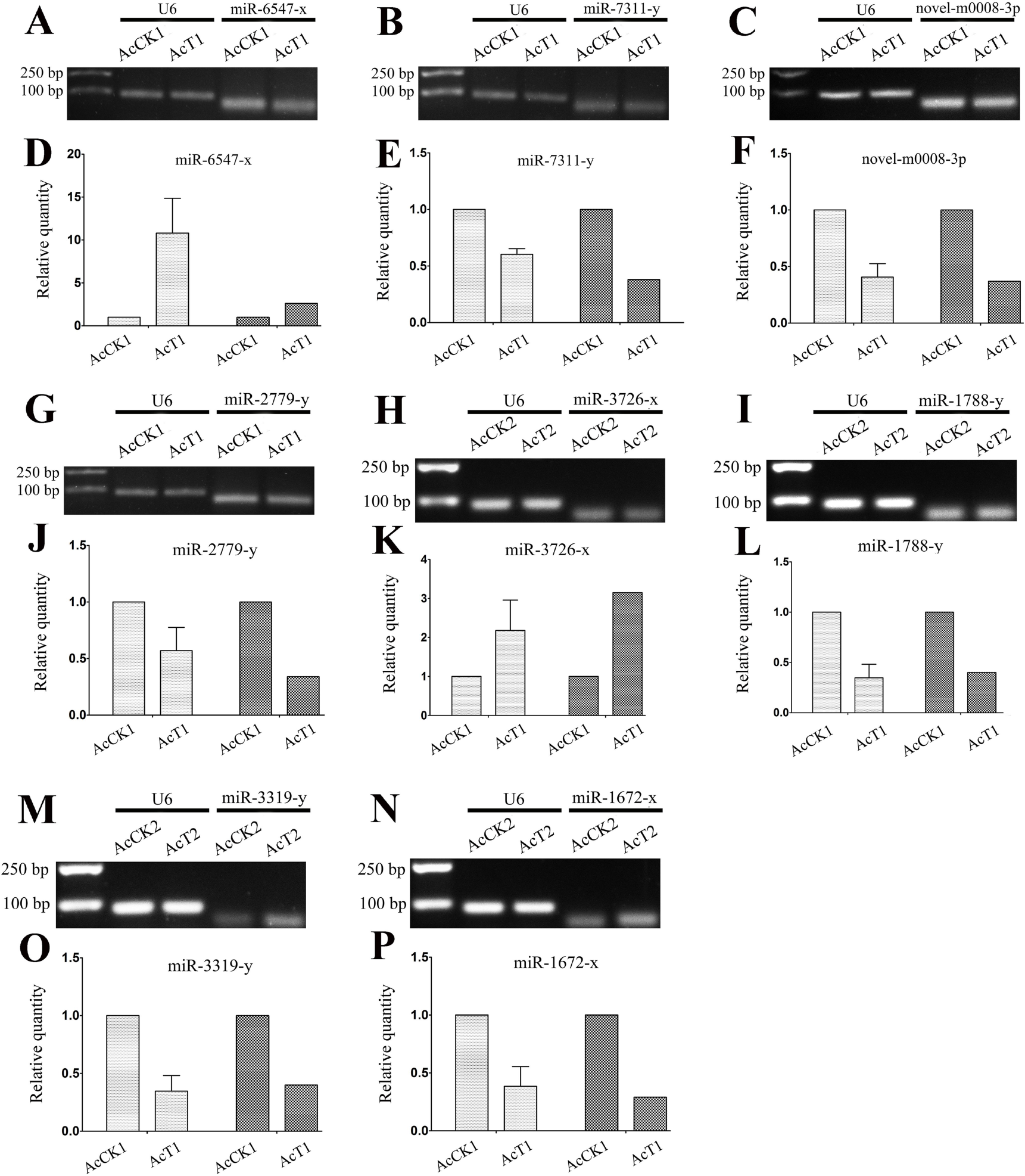
Stem-loop RT-PCR and RT-qPCR validation of *A. c. cerana* DEmiRNAs. (A-C) Stem-loop RT-PCR validation of miR-6547-x, miR-7311-x, and novel-m0008-3p; (G-I) Stem-loop RT-PCR validation of miR-2779-y, miR-3726-x, and miR-1788-y; (M-N) Stem-loop RT-PCR validation of miR-3319-y and miR-1672-x; (D-F) RT-qPCR validation of miR-6547-x, miR-7311-x, and novel-m0008-3p; (J-L) RT-qPCR validation of miR-2779-y, miR-3726-x, and miR-1788-y; (O-P) RT-qPCR validation of miR-3319-y and miR-1672-x; snRNA U6 was used as a positive control in stem-loop RT-PCR and reference gene in RT-qPCR.

## 4. Discussion

Previous studies have mainly been focused on western honeybee miRNAs^50–54^. However, relevant studies on the eastern honeybee were extremely limited. Shi et al. previously performed Illumina sequencing of miRNAs in the royal jelly from *A. mellifera* and *A. cerana*, and the results showed that there were differences between them; they further found that the transcriptomes of *A. mellifera* adults were affected by the two kinds of royal jelly^55^. Over the years, increasing evidence has demonstrated that miRNAs play vital roles in host-pathogen interactions via regulation of the gene expression of hosts or pathogens^56–58^. Although advances have been made regarding the actions of miRNAs in the interactions between the western honeybee and *N. ceranae*^59^, the miRNA profiles and DEmiRNAs in eastern honeybee-*N. ceranae* interactions remains completely unknown. In a previous study, Huang et al. deep-sequenced *A. mellifera* miRNAs daily across the six-day reproductive cycle of *N. ceranae* and identified 17 DEmiRNAs that target more than 400 genes^25^. Here, in order to identify miRNAs involved in the eastern honeybee responding to *N. ceranae* infection, we constructed for the first time four small RNA libraries from untreated and *N. ceranae*-treated *A. c. ceranae* workers’ midgut samples. In total, 529 conserved and 25 novel miRNAs were identified from sRNA-seq data using bioinformatics, which enriches the miRNA reservoir of the eastern honeybee and provides candidate miRNAs for functional studies in the future. In addition, eight predicted miRNAs were confirmed to be expressed in the midguts of *A. c. cerana* workers via stem-loop RT-PCR (**Figure 4**). Additionally, we also validated the expression of another eight DEmiRNAs (**Figure 9A-C, G-I, M-N**). Deep sequencing has become the mainstream method to explore novel miRNAs at present^60^. Notably, all of the 25 novel miRNAs predicted in this work were weakly expressed, in accordance with the results for some lower invertebrates such as *Drosophila* and *Cyprinus carpio*^49,61^. In our study, the most abundant miRNAs in AcCK1 and AcCK2 were bantam-y, miR-184-y, miR-1-y, miR-276-y, miR-750-y, miR-3477-x, miR-283-x, miR-12-z, miR-31-x and miR-9-z with TPM ≥ 31,625.97 each (**Table 3**). Surprisingly, these miRNAs were also highly expressed in AcT1 and AcT2 (**Table 3**). As a versatile player in many biological processes in *Drosophila*, *bantam* regulates fly development^62–63^, cell proliferation and apoptosis in many cell types^64–65^, and the production of the molting hormone ecdysone^66^. Through systematic manipulation of Dpp signaling in the *Drosophila* wing, Zhang et al. discovered that Dpp promoted proliferation in the lateral wing disc and repressed proliferation in the medial wing disc via *omb*, which controlled the regional proliferation rate by oppositely regulating transcription of *bantam* in the medial versus lateral wing disc^62^. Wu et al. previously investigated the role of the *bantam* miRNA in the regulation of neuroblast homeostasis in the *Drosophila* brain and found that *bantam* is a direct transcriptional target of the Notch signaling pathway; in addition, bantam feedback regulates the pathway by negatively regulating its target mRNA *numb*^63^. In the current work, *bantam*-y was the miRNA with the highest expression level in both AcCK1 and AcCK2 (**Table 3**), implying its key role in the regulation of the development of the *A. c. cerana* workers’ midguts. MiR-2, an invertebrate-specific miRNA family that has been predicted in fruit flies, is highly expressed in the heads of *Drosophila* and *Bombyx mori*^67–68^ and the neurons in *Caenorhabditis*^69^. It has also been shown to participate in the regulation of apoptosis of neuroblasts during the normal development of the nervous system in *Drosophila*^70^. Recent evidence shows that miR-2 regulates functions essential for the normal wing morphogenesis of *B. mori* via targeting of *awd* and *fng*^71^. Alexandra’s group revealed that miRs-2/13, including miR-2a, miR-2b, miR-13a, and miR-13b, play an important regulatory part in the embryonic development of *Drosophila* by targeting nine genes, including *Sos* and *Myd88*^72^. In the present study, miR-2-y and miR-13-y were detected to be highly expressed in the normal midguts of *A. c. cerana* workers (**Table 3**), which suggests these 2 miRNAs may play a fundamental role in regulating the development and morphogenesis of the midgut and apoptosis of the midgut cells. However, the functions of other miRNAs that were abundant in *A. c. cerana* midgut samples, such as miR-750-y and miR-3477-x, remain largely unknown.

DEmiRNAs in the midguts of *A. c. cerana* workers challenged by *N. ceranae* are particularly worthy of attention since they are likely to be direct regulators of host-pathogen interactions. In this research, 14 and 12 DEmiRNAs were identified in AcT1 VS AcCK1 and AcT2 VS AcCK2, respectively. Among them, miR-60-y, miR-8462-x, miR-2965-y, miR-676-y, and miR-6313-y were shared by the two comparison groups, implying their primary roles in host responses to *N. ceranae* infections. In addition, nine DEmiRNAs, including miR-1-x, miR-980-y, miR-965-x, miR-598-y, miR-6717-x, miR-4635-y, miR-9204-x, miR-3726-x, and miR-252-y, were specific in AcT1 VS AcCK1, and all of them were conserved miRNAs (**Figure 5B**, see also **Table S2** and **Table S3**). Additionally, seven DEmiRNAs, including two novel miRNAs (novel-m0003-3p and novel-m0019-5p), were specific in AcT2 VS AcCK2 (**Figure 5B**, see also **Table S2** and **Table S3**). We believe these unique DEmiRNAs play specific parts during different stages of host responses under the stress of *N. ceranae*.

To gain further insight into the potential genes associated with *N. ceranae* infection, we conducted GO analysis for DEmiRNA target genes to identify biologically important terms. The mostly enriched terms for the target genes in AcT1 VS AcCK1 and AcT2 VS AcCK2 were binding, cellular process, biological regulation, single-organism process, and catalytic activity (**Figure 6**, see also **Table S4** and **Table S5**), indicating the regulation of host metabolism and cellular activity via DEmiRNAs in response to *N. ceranae*. Huang and colleagues found that the target genes of DEmiRNAs in the *N. ceranae*-infected midgut of the *A. mellifera* workers were mostly involved in binding, signaling, nucleus, transmembrane transport, and DNA binding, which differ from our findings in the present study. This difference reflects the different responses of the two honeybee species with different resistances to the same fungal pathogen, *N. ceranae*. Considering the fact that *A. cerana* is more resistant to *N. ceranae* compared to *A. mellifera*, we inferred that DEmiRNAs may in part play a special role in the *N. ceranae*-resistance difference. In addition, we found 88 and 89 DEmiRNA-targeted genes in AcT1 VS AcCK1 and AcT2 VS AcCK2 were respectively enriched in response to stimulus, and there was one target gene for DEmiRNAs in AcT1 VS AcCK1 enriched in immune system process, which indicates the regulatory roles of the corresponding DEmiRNAs in host immunity defenses against *N. ceranae*. Furthermore, KEGG analysis for the target genes was performed to investigate the pathways influenced by *N. ceranae*. The result showed that out of the 104 pathways enriched by target genes of DEmiRNAs in AcT1 VS AcCK1, 62 were related to material and energy metabolism, such as carbohydrate metabolism including the pentose phosphate pathway and citrate cycle, lipid metabolism including arachidonic acid metabolism and fatty acid biosynthesis, nucleotide metabolism including purine metabolism and pyrimidine metabolism, energy metabolism including sulfur metabolism and oxidative phosphorylation (**Figure 7A**, see also **Table S6**). A total of 92 pathways were enriched by DEmiRNA target genes in AcT2 VS AcCK2; among them, 52 pathways were associated with material and energy metabolism (**Figure 7B**, see also **Table S7**). These results demonstrated that host DEmiRNAs regulate metabolism-related genes in response to *N. ceranae* invasion. Meanwhile, immunity-related pathways were further analyzed, and the results showed that endocytosis, phagosome, lysosome, ubiquitin mediated proteolysis, Jak-STAT signaling pathway, and MAPK signaling pathways were enriched by the target genes in AcT1 VS AcCK1 (**Figure 7A**, see also **Table S6**). Surprisingly, these immune pathways were also enriched by DEmiRNA target genes in AcT2 VS AcCK2 (**Figure 7B**, see also **Table S7**). These results demonstrated that DEmiRNAs and their target genes were involved in cellular and humoral immune responses of the host to *N. ceranae* invasion. Together, these findings indicate that host material and energy metabolism, cellular activity, and immunity defenses were significantly impacted by *N. ceranae* infection, and DEmiRNAs played a comprehensive role in host-pathogen interactions.

MiR-1 is a muscle-specific miRNA that plays important roles in regulating heart development and muscle differentiation^73–74^. It has also been known to be a tumor suppressor gene that not only inhibits cancer cell proliferation, metastasis, and invasion but also enhances apoptosis by regulating oncogenic targets in many cancer cells^75–76^. In insects, miR-1 was found to be involved in immune processes in *Drosophila melanogaster*^77^, and it was differentially expressed during pathogen infection in *Aedes aegypti*^78^. In this work, miR-1-x was dramatically down-regulated in AcT1 VS AcCK1, suggesting that the host and fungal pathogen can interact with each other via miR-1-x by regulating host proliferation and apoptosis and immune response. Wu et al. carried out Illumina sequencing of dengue virus-2 (DENV-2)-infected and uninfected *Aedes albopictus* and observed a series of DEmiRNAs including miR-252^79^. By using RT-qPCR, Yan et al. detected a high expression level of miR-252 in DENV-2-infected C6/36 cells and mosquitoes compared with uninfected cells and mosquitos and further revealed that DENV-2 envelope protein expression can be affected by the overexpression of miR-252 with a mimic or down-regulation using an inhibitor^80^. Recently, Lim and colleagues discovered that miR-252-5p can control the cell cycle by directly repressing Abelson interacting protein (Abi) in *Drosophila* S2 cells by using cross-linking immunoprecipitation and deep sequencing of endogenous Argonaute 1 (Ago1) protein^81^. In our study, miR-252-y had a weak up-regulation in the midgut 7 dpi with *N. ceranae*, suggesting that miR-252-y may participate in *N. ceranae* defense in *A. c. cerana*. As a member of the miR-17-92 cluster, miR-92a inhibits apoptosis and promotes the proliferation of other cell types, including endothelial cells^82–83^. Apoptosis is of great importance for development and pathogen defense in multicellular organisms including insects^84–85^. Previous studies have demonstrated that *Spodoptera frugiperda* larvae and *B. mori* larvae battle baculovirus infection by selective apoptosis of infected cells from the midgut epithelium^86–87^. *N. ceranae* is able to enhance its development during the infection process by preventing apoptosis in epithelial cells of infected *A. mellifera*^88^. A previous study showed that miR-194 expression was markedly decreased in A549 alveolar epithelial cells following infection with influenza A virus (IAV), and miR-194 could suppress fibroblast growth factor 2 (FGF2) expression at the mRNA and protein levels^89^. Xie and colleagues suggested that miR-194 increases cell apoptosis by inhibiting the NF-*κ*B pathway in WI38 cells, and it may be used as a potential targeted therapy for the treatment of infantile pneumonia^90^. In this present study, miR-92-x had a sharp down-regulation, while miR-194-y was significantly up-regulated in the *A. c. cerana* midgut 10 dpi with *N. ceranae*. Therefore, we inferred that *A. c. cerana* could fight *N. ceranae* by promoting apoptosis of host midgut cells via regulation of the expression levels of miR-92-x and miR-194-y. Further studies are needed to confirm the mechanism underlying the roles of above-mentioned DEmiRNAs in host-pathogen interactions.

A single miRNA can regulate several target genes at the same time to inhibit their expression and vice versa^91^. miR-598 has been found to play inhibitory role in osteosarcoma progression *in vivo* and *in vitro* by modulating osteoblastic differentiation in the microenvironment and targeting *PDGFB* and *MET*. Based on expressed sequence tags (ESTs) resources from the LNCaP cells, Saravanan et al. identified an hsa-miR-3654 with a higher expression level in LNCaP cells than the normal and androgen insensitive prostate cancer cell lines (PNT1A, PC-3)^92^. To our knowledge, there is no documentation of miR-598 and miR-3654 in insects until now. In the current work, miR-598 lies in the center of the regulation networks of DEmiRNAs in AcT1 VS AcCK1, linking to as many as 45 target genes such as XM_017048388.1, XM_017049101.1, and XM_017051459.1 (**Figure 8A**), while miR-3654-y was a key miRNA in the regulation networks of DEmiRNAs in AcT2 VS AcCK2, binding to four targets including XM_017057343.1, XM_017057344.1, XM_017057345.1, and XM_017057346.1 (**Figure 8B**). These results indicated that miR-598 and miR-3654-y are likely to be key regulators of the interactions between *A. c. cerana* and *N. ceranae*, and these two miRNAs would be our candidates for future studies of host immune response and resistance to *N. ceranae*.

In conclusion, using high-throughput sequencing technology and bioinformatics, we systematically investigated the highly expressed miRNAs in the normal *A. c. cerana* midgut, and DEmiRNAs as well as their target genes in the *N. ceranae*-infected midgut. This study provides the first global view of the miRNA profiles and DEmiRNAs in the midgut of the *A. c. cerana* worker invaded by *N. ceranae*. Findings in the present study demonstrated that *N. ceranae* invasion causes alterations in the expression of miRNAs that regulate metabolism, cellular activity, and immune response in the host. Taken together, our results not only provide novel insights into understanding *A. c. cerana*-*N. ceranae* interactions but also lay a foundation for deciphering the molecular mechanisms underlying the resistance of the eastern honeybee to *N. ceranae*.

## Author Contributions

RG designed this study. DF-C, YD, HZ-C, HP-W, CL-X, YZ-Z and CS-H carried out laboratory work. RG, DF-C, YD and QY-D performed bioinformatic analysis. RG, DF-C and YD supervised the work and contributed to preparation of the manuscript.

## Acknowledgements

This research was supported by the Earmarked Fund for China Agriculture Research System (CARS-44-KXJ7), the Science and Technology Planning Project of Fujian Province (2018J05042), the Teaching and Scientific Research Fund of Education Department of Fujian Province (JAT170158), the Outstanding Scientific Research Manpower Fund of Fujian Agriculture and Forestry University (xjq201814), the Scientific and Technical Innovation Fund of Fujian Agriculture and Forestry University (CXZX2017343, CXZX2017343). Rui Guo is very grateful to his wife Qian Cai for continuous assistance and great love.

---

**Figure S1.**
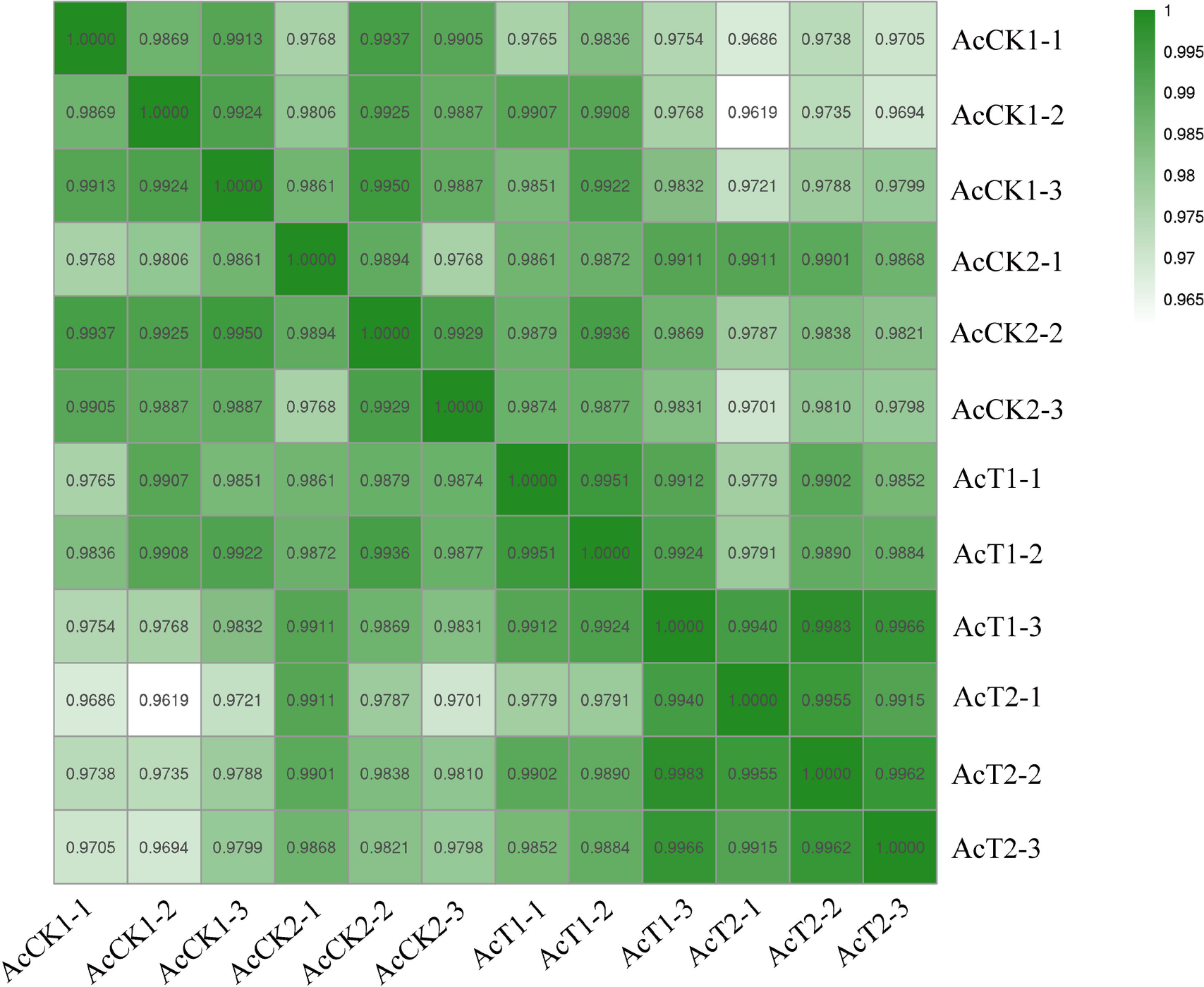
Pearson correlation between different biological repeats within each gut sample from workers’ midgut of *Apis cerana cerana*

## References

1. Bromenshenk, J.J., Henderson, C.B., Seccomb, R.A., Welch, P.M., Debnam, S.E., and Firth, D.R. 2015. Bees as biosensors: chemosensory ability, honey bee monitoring systems, and emergent sensor technologies derived from the pollinator syndrome. Biosensors, 5, 678–711.

2. Hu, X., Ke, L., Wang, Z., and Zeng, Z. 2018. Dynamic transcriptome landscape of Asian domestic honeybee (*Apis cerana*) embryonic development revealed by high-quality RNA sequencing. BMC Dev. Biol., 18, 11.

3. Elsik, C.G., Worley, K.C., Bennett, A.K., Beye, M., Camara, F., Childers, C.P., et al. 2014. Finding the missing honey bee genes: lessons learned from a genome upgrade. BMC Genomics, 15, 86.

4. Park, D., Jung, J.W., Choi, B.S., Jayakodi, M., Lee, J., Lim, J., et al. 2014. Uncovering the novel characteristics of Asian honey bee, *Apis cerana*, by whole genome sequencing. BMC Genomics, 16, 1–16.

5. Oldroyd, B.P., and Wongsiri, S. 2006. Asian Honey Bees: Biology, Conservation, and Human Interactions. Cambridge: Harvard University Press.

6. Liu, Z., Yao, P., Guo, X., and Xu, B. 2014. Two small heat shock protein genes in *Apis cerana cerana*: characterization, regulation, and developmental expression. Gene, 545, 205–214.

7. Wittner, M., and Weiss, L.M. 1999. The Microsporidia and Microsporidiosis. Washington, DC: ASM Press.

8. Visvesvara, G.S. 2002. *In vitro* cultivation of microsporidia of clinical importance. Clin. Microbiol. Rev., 15, 401–413.

9. Chen, Y.P., Evans, J.D., Murphy, C., Gutell, R., Zuker, M., Gundensen-Rindal, D., et al. 2010. Morphological, molecular, and phylogenetic characterization of *Nosema ceranae*, a microsporidian parasite isolated from the European honey bee, *Apis mellifera*. J. Eukaryot. Microbi., 56, 142–147.

10. Adl, S.M., Simpson, A.G.B., Farmer, M.A., Andersen, R.A., Anderson, O.R., Barta, J.R., et al. 2005. The new higher level classification of eukaryotes with emphasis on the taxonomy of protists. J. Eukaryot. Microbiol., 52, 399–451.

11. Fries, I., Feng, F., daSilva, A., Slemenda, S.B., and Pieniazek, N.J. 1996. *Nosema ceranae* nsp (Microspora,Nosematidae), morphological and molecular characterization of a microsporidian parasite of the Asian honey bee *Apis cerana* (Hymenoptera,Apidae). Europ. J. Protistol., 32, 356–365.

12. Higes, M., Martin-Hernandez, R., and Meana, A. 2006. *Nosema ceranae*, a new microsporidian parasite in honeybees in Europe. J. Invertebr. Pathol., 92, 93–95.

13. Huang, W.F., Jiang, J.H., Chen, Y.W., and Wang, C.H. 2007. A *Nosema ceranae* isolate from the honeybee *Apis mellifera*. Apidologie, 38, 30–37.

14. Klee J, Besana A M, Genersch E, et al. 2007. Widespread dispersal of the microsporidian *Nosema ceranae*, an emergent pathogen of the western honey bee, *Apis mellifera*. J. Invertebr. Pathol., 96, 1–10.

15. Chen, Y., Evans, J.D., Smith, I.B., and Pettis, J.S. 2008. *Nosema ceranae* is a long-present and wide-spread microsporidian infection of the European honey bee (*Apis mellifera*) in the United States. J. Invertebr. Pathol., 97, 186–188.

16. Alaux, C., Brunet, J.L., Dussaubat, C., Mondet, F., Tchamitchan, S., Cousin, M., et al. 2010. Interactions between Nosema microspores and a neonicotinoid weaken honeybees (*Apis mellifera*). Environ. Microbiol., 12, 774–782.

17. Pettis, J.S., Vanengelsdorp, D., Johnson, J., and Dively, G. 2012. Pesticide exposure in honey bees results in increased levels of the gut pathogen *Nosema*. Naturwissenschaften, 99, 153–158.

18. Bartel, D.P. 2004. MicroRNAs: genomics, biogenesis, mechanism, and function. Cell, 116, 281–297.

19. Liu, J., Wang, X., Yang, X., Liu, Y., Shi, Y., Ren, J., et al. 2014. miRNA423-5p regulates cell proliferation and invasion by targeting trefoil factor 1 in gastric cancer cells. Cancer Lett., 347, 98–104.

20. Hammond, S.M. 2006. MicroRNAs as oncogenes. Curr. Opin. Genet. Dev. 16, 4.

21. Cristino, A.S., Tanaka, E.D., Rubio, M., Piulachs, M.D., and Belles, X. 2011. Deep sequencing of organ- and stage-specific microRNAs in the evolutionarily basal insect *Blattella germanica* (L.) (Dictyoptera, Blattellidae). PLoS One, 6, e19350.

22. Wagschal, A., Najafishoushtari, S.H., Wang, L., Goedeke, L., Sinha, S., deLemos, A.S. et al. 2015. Genome-wide identification of microRNAs regulating cholesterol and triglyceride homeostasis. Nat. Med., 21, 1290–1297.

23. Leung, A.K.L., and Sharp, P.A. 2010. MicroRNA functions in stress responses. Mol. Cell, 40, 280–293.

24. Ma, F., Xu, S., Liu, X., Zhang, Q., Xu, X., Liu, M., et al. 2011. The microRNA miR-29 controls innate and adaptive immune responses to intracellular bacterial infection by targeting interferon-γ. Nat. Immunol., 12, 861–869.

25. Huang, Q., Chen, Y., Rui. W.W., Schwarz, R.S., and Evans, J.D. 2015. Honey bee microRNAs respond to infection by the microsporidian parasite *Nosema ceranae*. Sci. Rep., 5, 17494.

26. Higes, M., Garcíapalencia, P., Martínhernández, R., and Meana, A. 2007. Experimental infection of *Apis mellifera* honeybees with *Nosema ceranae* (Microsporidia). J. Invertebr. Pathol., 94, 211–217.

27. Cornman, R.S., Chen, Y.P., Schatz, M.C., Street, C., Zhao, Y., Desany, B., et al. 2009. Genomic analyses of the microsporidian *Nosema ceranae*, an emergent pathogen of honey bees. PLoS Pathog., 5, e1000466.

28. Fries, I., Chauzat, M.P., Chen, Y.P., Doublet, V., Genersch, E., Gisder, S., et al. 2013. Standard methods for *Nosema* research. J. Apicult. Res., 52, 1–28.

29. Forsgren, E., and Fries, I. 2010. Comparative virulence of *Nosema ceranae* and *Nosema apis* in individual European honey bees. Vet. Parasitol., 170, 212–217.

30. Langmead, B., Trapnell, C., Pop, M., and Salzberg, S.L. 2009. Ultrafast and memory-efficient alignment of short DNA sequences to the human genome. Genome Biol., 10, R25.

31. Kozomara, A., Griffiths-Jones, S. 2010. miRBase: integrating microRNA annotation and deep-sequencing data. Nucleic acids Res., 39, D152–D157.

32. Hofacker, I.L. 2009. RNA secondary structure analysis using the Vienna RNA package. Curr. Protoc. Bioinforma., Chapter 12, Unit12.2.

33. Zhu, E., Zhao, F., Xu, G., Hou, H., Zhou, L., Li, X., et al., 2010. mirTools: microRNA profiling and discovery based on high-throughput sequencing. Nucleic acids Res., 38, W392–W397.

34. Betel, D., Wilson, M., Gabow, A., Marks, D.S., Sander, C. 2008. The microRNA. org resource: targets and expression. Nucleic Acids Res., 36, D149–D153.

35. Rehmsmeier, M., Steffen, P., Höchsmann, M., and Giegerich, R. 2004. Fast and effective prediction of microRNA/target duplexes. RNA, 10, 1507–1517.

36. Allen, E., Xie, Z., Gustafson, A.M., and Carrington, J.C. 2005. microRNA-directed phasing during trans-acting siRNA biogenesis in plants. Cell, 121, 207–221.

37. Smoot, M.E., Ono, K., Ruscheinski, J., Wang, P.L., and Ideker, T. 2011. Cytoscape 2.8: new features for data integration and network visualization. Bioinformatics, 27, 431–432.

38. Conesa, A., Götz, S., García-Gómez, J.M., Terol, J., Talón, M., and Robles, M. 2005. Blast2GO: a universal tool for annotation, visualization and analysis in functional genomics research. Bioinformatics, 21, 3674–3676.

39. Ye, J., Fang, L., Zheng, H., Zhang, Y., Chen, J., Zhang, Z., et al. 2006. WEGO: a web tool for plotting GO annotations. Nucleic Acids Res., 34, W293–W297.

40. Kanehisa, M., Araki, M., Goto, S., Hattori, M., Hirakawa, M., Itoh, M., et al. 2008. KEGG for linking genomes to life and the environment. Nucleic Acids Res., 36, D480–D484.

41. Chen, C.F., Ridzon, D.A., Broomer, A.J., Zhou, Z.H., Lee, D.H., Nguyen, J.T., et al. 2005. Real-time quantification of microRNAs by stem-loop RT-PCR. Nucleic Acids Res., 33, e179.

42. Livak, K.J., and Schmittgen, T.D. 2001. Anaysis of relative gene expression data using realtime quantitative PCR and the 2^−ΔΔCt^ method. Methods., 25, 402–408.

43. Kudla, G., Lipinski, L., Caffin, F., Helwak, A., and Zylicz, M. 2006. High guanine and cytosine content increases mRNA levels in mammalian cells. PLoS Biol., 4, e180.

44. Shepotinovskaya, I.V., and Uhlenbeck, O.C. 2008. Catalytic diversity of extended hammerhead ribozymes. Biochemistry, 47, 7034–7042.

45. Lewis, B.P., Burge, C.B., and Bartel, D.P. 2005. Conserved seed pairing, often flanked by adenosines, indicates that thousands of human genes are microRNA targets. Cell, 120, 15–20.

46. Dezulian, T., Palatnik, J.F., Huson, D., and Weigel, D. 2005. Conservation and divergence of microRNA families in plants. Genome Biol., 6, 1–25.

47. Ai, L., Xu, M.J., Chen, M.X., Zhang, Y.N., Chen, S.H., Guo, J., et al. 2012. Characterization of microRNAs in *Taenia saginata* of zoonotic significance by Solexa deep sequencing and bioinformatics analysis. Parasitol. Res., 110, 2373–2378.

48. Ji, Z., Wang, G., Xie, Z., Zhang, C., and Wang, J. 2012. Identification and characterization of microRNA in the dairy goat (*Capra hircus*) mammary gland by Solexa deep-sequencing technology. Mol. Biol. Rep., 39, 9361–9371.

49. Wang, F., Jia, Y., Wang, P., Yang, Q., Du, Q., and Chang, Z. 2017. Identification and profiling of *Cyprinus carpio* microRNAs during ovary differentiation by deep sequencing. BMC Genomics, 18, 333–349.

50. Chen, X., Yu, X., Cai, Y., Zheng, H., Yu, D., Liu, G., et al. 2010. Next-generation small RNA sequencing for microRNAs profiling in the honey bee *Apis mellifera*. Insect Mol. Biol., 19, 799–805.

51. Liu, F., Peng, W., Li, Z., Li, W., Li, L., Pan, J., et al. 2012. Next-generation small RNA sequencing for microRNAs profiling in *Apis mellifera*: comparison between nurses and foragers. Insect Mol. Biol., 21, 297–303.

52. Zondag, L., Dearden, P.K., and Wilson, M.J. 2012. Deep sequencing and expression of microRNAs from early honeybee (*Apis mellifera*) embryos reveals a role in regulating early embryonic patterning. BMC Evol. Biol., 12, 211–223.

53. Qin, Q.H., Wang, Z.L., Tian, L.Q., Gan, H.Y., Zhang, S.W., and Zeng, Z.J. 2014. The integrative analysis of microRNA and mRNA expression in *Apis mellifera*, following maze-based visual pattern learning. Insect Sci., 21, 619–636.

54. Ashby, R., Forêt, S., Searle, I., and Maleszka, R. 2016. MicroRNAs in honey bee caste determination. Sci. Rep., 6:18794–18808.

55. Shi, Y.Y., Wu, X.B., Huang, Z.Y., Wang, Z.L., Yan, W.Y., and Zeng, Z.J. 2012. Epigenetic modification of gene expression in honey bees by heterospecific gland secretions. PLoS One, 7, e43727.

56. Tu, Z., and Edward, M. 2008. Cloning, characterization, and expression of microRNAs from the Asian malaria mosquito, *Anopheles stephensi*. BMC Genomics, 9, 244.

57. Padmanabhan, C., Zhang, X., and Jin, H. 2009. Host small RNAs are big contributors to plant innate immunity. Curr. Opin. Plant Biol., 12, 465–472.

58. Singh, C.P., Singh, J., and Nagaraju, J. 2012. A baculovirus-encoded microRNA (miRNA) suppresses its host miRNA biogenesis by regulating the exportin-5 cofactor Ran. J. Virol., 86, 7867–7879.

59. Huang, Q., and Evans, J.D. 2015. Identification of microRNA-like small RNAs from fungal parasite *Nosema ceranae*. J. Inverteb. Pathol., 133, 107–109.

60. Wojcicka, A., Swierniak, M., Kornasiewicz, O., Gierlikowski, W., Maciag, M., Kolanowska, M., et al. 2014. Next generation sequencing reveals microRNA isoforms in liver cirrhosis and hepatocellular carcinoma. Int. J. Biochem. Cell Biol., 53, 208–217.

61. Lu, J., Shen, Y., Wu, Q., Kumar, S., He, B., Shi, S., et al. 2008. The birth and death of microRNA genes in *Drosophila*. Nat. Genet., 40, 351–355.

62. Zhang, X., Luo, D., Pflugfelder, G.O., and Shen, J. 2013. Dpp signaling inhibits proliferation in the *Drosophila* wing by Omb-dependent regional control of *bantam*. Development, 140, 2917–2922.

63. Wu, Y.C., Lee, K.S., Song, Y., Gehrke, S., and Lu, B. 2017. The bantam microRNA acts through *Numb* to exert cell growth control and feedback regulation of Notch in tumor-forming stem cells in the *Drosophila* brain. PLoS Genet., 13, e1006785.

64. Dong, L., Li, J., Huang, H., Yin, M.X., Xu, J., Li, P.., et al. 2015. Growth suppressor lingerer regulates *bantam* microRNA to restrict organ size. J Mol. Cell Biol., 7, 415–428.

65. Catarina, B.P., Fernando, C., and Florence, J. 2015. The retinal determination gene *Dachshund* restricts cell proliferation by limiting the activity of the Homothorax-Yorkie complex. Development, 142, 1470–1479.

66. Boulan, L., Martín, D., and Milán, M. 2013. *bantam* miRNA promotes systemic growth by connecting insulin signaling and ecdysone production. Current Biol., 23, 473–478.

67. Liu, N., Okamura, K., Tyler, D.M., Phillips, M.D., Chung, W.J., and Lai, E.C. 2008. The evolution and functional diversification of animal microRNA genes. Cell Res., 18, 985–996.

68. Xiang, Z., Yin, J., Zhang, D., Yin, J., Xiang, Z., and Xia, Q. 2010. MicroRNAs show diverse and dynamic expression patterns in multiple tissues of *Bombyx mori*. BMC Genomics, 11, 85–96.

69. Martinez, N.J., Ow, M.C., Reece-Hoyes, J.S., Barrasa, M.I., Ambros, V.R., and Walhout, A.J. 2008. Genome-scale spatiotemporal analysis of *Caenorhabditis elegans* microRNA promoter activity. Genome Res., 18, 2005–2015.

70. Tan, Y., Yamada-Mabuchi, M., Arya, R., St Pierre, S., Tang, W., Tosa, M., et al. 2011. Coordinated expression of cell death genes regulates neuroblast apoptosis. Development, 138, 2197–2206.

71. Ling, L., Ge, X., Li, Z., Zeng, B., Xu, J., Chen, X., et al. 2015. MiR-2 family targets awd and fng to regulate wing morphogenesis in *Bombyx mori*. RNA Biol., 12, 742–748.

72. Boutla, A., Delidakis, C., and Tabler, M. 2003. Developmental defects by antisense-mediated inactivation of microRNAs 2 and 13 in *Drosophila* and the identification of putative target genes. Nucleic Acids Res., 31, 4973–4980.

73. Kwon, C., Han, Z., Olson, E.N., et al. 2005. MicroRNA1 influences cardiac differentiation in *Drosophila* and regulates Notch signaling. Proc. Natl. Acad. Sci. USA., 102, 18986–18991.

74. Sokol, N.S., and V. Ambros. 2005. Mesodermally expressed Drosophila microRNA-1 is regulated by Twist and is required in muscles during larval growth. Genes Dev., 19, 2343–2354.

75. Tang, Y., Zheng, J., Sun, Y., Wu, Z., Liu, Z., and Huang, G. 2009. MicroRNA-1 regulates cardiomyocyte apoptosis by targeting *Bcl-2*. Int. Heart J., 50, 377–387.

76. Yan, H.B., Huang, J.C., Chen, Y.R., Yao, J.N., Cen, W.N., Li, J.Y., et al. 2018. Role of miR-1 expression in clear cell renal cell carcinoma (ccRCC): A bioinformatics study based on GEO, ArrayExpress microarrays and TCGA database. Pathol. Res. Pract., 214, 195–206.

77. Fullaondo, A., and Lee, S.Y. 2012. Identification of putative miRNA involved in *Drosophila melanogaster* immune response. Dev. Comp. Immunol., 36, 267–273.

78. Shrinet, J., Jain, S., Jain, J., Bhatnagar, R.K., and Sunil, S. 2014. Next generation sequencing reveals regulation of distinct *Aedes* microRNAs during chikungunya virus development. PLoS Negl. Trop. Dis., 8, e2616.

79. Wu, J.Y., Zheng, P.M., Tu, Z.J., and Chen, X.J. 2010. OL-032 identification of miRNAs in *Aedes albopictus*, and determination of their expression profiles during developmental stages and blood feeding using Solexa sequencing. Int. J. Infect. Dis., 14, S14–S15.

80. Yan, H., Zhou, Y., Liu, Y., Deng, Y., and Chen, X. 2014. miR-252 of the Asian tiger mosquito *Aedes albopictus* regulates dengue virus replication by suppressing the expression of the dengue virus envelope protein. J. Med. Virol., 86, 1428–1436.

81. Lim, D.H., Lee, S., Han, J.Y., Choi, M.S., Hong, J.S., Seong, Y., et al. 2018. Ecdysone-responsive microRNA-252-5p controls the cell cycle by targeting *Abi* in *Drosophila*. FASEB J., 32, 4519–4533.

82. Liu, J., Yao, W., Yao, Y., Du, X., Zhou, J., Ma, B., et al. 2014. MiR-92a inhibits porcine ovarian granulosa cell apoptosis by targeting *Smad7* gene. FEBS Lett., 588, 4497–4503.

83. Zhang, L., Zhou, M., Qin, G., Weintraub, NL., and Tang, Y. 2014. MiR-92a regulates viability and angiogenesis of endothelial cells under oxidative stress. Biochem. Biophys. Res. Commun., 446, 952–958.

84. Kurze, C., Le, C.Y., Dussaubat, C., Erler, S., Kryger, P., Lewkowski, O., et al. 2015. Nosema tolerant honeybees (*Apis mellifera*) escape parasitic manipulation of apoptosis. PLoS One, 10, e0140174.

85. Guo, R., Wang, S., Xue, R., Cao, G., Hu, X., Huang, M., et al. 2015. The gene expression profile of resistant and susceptible *Bombyx mori* strains reveals cypovirus-associated variations in host gene transcript levels. Appl. Microbiol. Biotechnol., 99, 5175–5187.

86. Clarke, T.E., and Clem, R.J. 2002. Lack of involvement of haemocytes in the establishment and spread of infection in *Spodoptera frugiperda* larvae infected with the baculovirus *Autographa californica* M nucleopolyhedrovirus by intrahaemocoelic injection. J. Gen. Virol., 83, 1565–1572.

87. Yao, H.P., Wu, X.F., and Gokulamma, K. 2006. Antiviral activity in the mulberry silkworm, *Bombyx mori* L. J. Zhejiang Univ. SCIENCE A, 7, 350–356.

88. Higes, M., Juarranz, Á., Dias-Almeida, J., Lucena, S., Botías, C., Meana, A., et al. 2013. Apoptosis in the pathogenesis of *Nosema ceranae* (Microsporidia:Nosematidae) in honey bees (*Apis mellifera*). Environ. Microbiol. Rep., 5, 530–536.

89. Wang, K., Lai, C., Gu, H., Zhao, L., Xia, M., Yang, P., et al. 2017. miR-194 inhibits innate antiviral immunity by targeting *FGF2* in influenza H1N1 virus infection. Front. Microbiol., 8, 2187–2196.

90. Xie, F., Yang, L., Han, L.L., and Yue, B. 2017. MicroRNA-194 regulates lipopolysaccharide-induced cell viability by inactivation of nuclear factor-*κ*B pathway. Cellular Physiol. Biochem., 43, 2470–2478.

91. Xie, W., Huang, A., Li, H., Feng, L.Z., Zhang, F.P., and Guo, W.S. 2016. Identification and comparative analysis of microRNAs in *Pinus massoniana*, infected by *Bursaphelenchus xylophilus*. Plant Growth Regul., 1–10.

92. Saravanan, S., Islam, V.I., Thirugnanasambantham, K., and Sekar, D. 2016. *In silico* identification of human miR 3654 and its targets revealed its involvement in prostate cancer progression. Microrna, 5(1): 1–6.

