## Supplemental Table 1 for "Comparative identification of microRNAs in *Apis cerana cerana* workers’ midguts responding to *Nosema ceranae* invasion"

**Table S1 Primers used in the present study.**

| **miRNA** | **Sequence** | |
| --- | --- | --- |
| miR-1943-x | Loop-1 | CTCAACTGGTGTCGTGGAGTCGGCAATTCAGTTGAGTGGTGCCC |
|  | F-1 | GGGAGGATCC |
| miR-3793-x | Loop-2 | CTCAACTGGTGTCGTGGAGTCGGCAATTCAGTTGAGCCGGCCAG |
|  | F-2 | AGCGTGTTTTCC |
| miR-252-y | Loop-3 | CTCAACTGGTGTCGTGGAGTCGGCAATTCAGTTGAGTGATAAGC |
|  | F-3 | CTGCTGCTCAAGT |
| miR-3963-x | Loop-4 | CTCAACTGGTGTCGTGGAGTCGGCAATTCAGTTGAGCGGTGTCA |
|  | F-4 | GTATCCCACTTC |
| miR-8516-x | Loop-5 | CTCAACTGGTGTCGTGGAGTCGGCAATTCAGTTGAGCTCCTGTT |
|  | F-5 | AGCGGCGAGCG |
| miR-149-y | Loop-6 | CTCAACTGGTGTCGTGGAGTCGGCAATTCAGTTGAGAGCCTCCC |
|  | F-6 | GAGGGAGGGA |
| miR-9008-x | Loop-7 | CTCAACTGGTGTCGTGGAGTCGGCAATTCAGTTGAGTTCGGGTC |
|  | F-7 | GCTGGCAGAA |
| novel-m0014-3p | Loop-8 | CTCAACTGGTGTCGTGGAGTCGGCAATTCAGTTGAGAGACTGCA |
|  | F-8 | CCGGCGATGATGTCA |
| miR-6547-x | Loop-9 | CTCAACTGGTGTCGTGGAGTCGGCAATTCAGTTGAGCTCCGTCC |
|  | F-9 | TGCGATGTGG |
| miR-7311-y | Loop-10 | CTCAACTGGTGTCGTGGAGTCGGCAATTCAGTTGAGTGGAGGCG |
|  | F-10 | CTCCGGGACG |
| miR-2779-y | Loop-11 | CTCAACTGGTGTCGTGGAGTCGGCAATTCAGTTGAGTGGTCCTT |
|  | F-11 | CGATCCGGCTCG |
| miR-3726-x | Loop-12 | CTCAACTGGTGTCGTGGAGTCGGCAATTCAGTTGAGAACGCTGG |
|  | F-12 | GAGTGGTGGATG |
| miR-1788-y | Loop-13 | CTCAACTGGTGTCGTGGAGTCGGCAATTCAGTTGAGCAGACTCG |
|  | F-13 | GGAGCGAAAG |
| miR-3319-y | Loop-14 | CTCAACTGGTGTCGTGGAGTCGGCAATTCAGTTGAGTTCACGGA |
|  | F-14 | CCTGCAACCCGGC |
| miR-1672-x | Loop-15 | CTCAACTGGTGTCGTGGAGTCGGCAATTCAGTTGAGCTCCCTTC |
|  | F-15 | GGTCAGGCCCG |
| novel-m0008-3p | Loop-16 | CTCAACTGGTGTCGTGGAGTCGGCAATTCAGTTGAGAAATGTCT |
|  | F-16 | CTCAATCCGTGTTG |
| U6 | F | GGCACTTTGTTAGGCTTTG |
|  | R | GTTTGGTGACTCTTCTTCCTTC |
|  | Universal-R | CTCAACTGGTGTCGTGGA |
