## Supplemental Table 2 for "Comparative identification of microRNAs in *Apis cerana cerana* workers’ midguts responding to *Nosema ceranae* invasion"

**Table S2 Summary of DEmiRNAs in AcT1 VS AcCK1 comparison group.**

| **miRNA** | **Sequence** | **Length** | **TPM of miRNAs**  **in AmCK1** | **TPM of miRNAs in AmT1** | **Log_2_FC** | ***P* value** | **Mark** |
| --- | --- | --- | --- | --- | --- | --- | --- |
| miR-676-y | GCTGTCCTAAGGTAGATGA | 19 | 0.01 | 80.42047 | 12.97335 | 2.96E-05 | Up |
| miR-60-y | ACATGTTCTGGTTGAAGA | 18 | 0.01 | 18.69587 | 10.8685 | 1.51E-05 | Up |
| miR-2965-y | AGGACTGCTACAGAGAGCA | 19 | 0.01 | 17.16527 | 10.74528 | 3.91E-05 | Up |
| miR-8462-x | ATTAATTTGATAAGTTATA | 19 | 0.01 | 14.42317 | 10.49417 | 0.000242 | Up |
| miR-6717-x | GCGATGTGGGGACGGAGA | 18 | 0.01 | 12.05733 | 10.2357 | 2.14E-07 | Up |
| miR-6313-y | TGCTGTGAAGTTTTGATT | 18 | 0.01 | 5.9736 | 9.222457 | 2.21E-06 | Up |
| miR-3726-x | GAGTGGTGGATGCCAGCGTT | 20 | 0.01 | 5.173933 | 9.015118 | 0.000162 | Up |
| miR-252-y | CTGCTGCTCAAGTGCTTATCA | 21 | 11.16776667 | 27.01673 | 1.274513 | 0.025931 | Up |
| miR-980-y | AAGCTGCCTTTTGAAGGGCAACA | 23 | 39.08906667 | 18.85337 | -1.05194 | 0.038812 | Down |
| miR-598-y | GTCGTCGTCGTCATCGTCA | 19 | 2.4139 | 0.01 | -7.91522 | 9.16E-05 | Down |
| miR-1-x | CCGTGCTTCCTTACTTCCCATA | 22 | 3.340533333 | 0.01 | -8.38393 | 2.82E-07 | Down |
| miR-965-x | GGGGAAAGGTTATAGCGATTATG | 23 | 3.6395 | 0.01 | -8.5076 | 4.12E-07 | Down |
| miR-4635-y | GAAGTCGGAACCCGCTAAG | 19 | 5.678066667 | 0.01 | -9.14926 | 4.11E-05 | Down |
| miR-9204-x | CTGGGATGAAATGTGGGT | 18 | 6.016033333 | 0.01 | -9.23267 | 0.00087 | Down |
