## Supplemental Table 3 for "Comparative identification of microRNAs in *Apis cerana cerana* workers’ midguts responding to *Nosema ceranae* invasion"

**Table S3 Summary of DEmiRNAs in AcT2 VS AcCK2 comparison group.**

| **miRNA** | **Sequence** | **Length** | **TPM of miRNAs**  **in AmCK2** | **TPM of miRNAs in AmT2** | **Log_2_FC** | ***P* value** | **Mark** |
| --- | --- | --- | --- | --- | --- | --- | --- |
| miR-676-y | GCTGTCCTAAGGTAGATGA | 19 | 0.01 | 410.3586 | 15.3246 | 4.02E-06 | Up |
| miR-60-y | ACATGTTCTGGTTGAAGA | 18 | 0.01 | 205.0298 | 14.32355 | 0.000287 | Up |
| miR-194-y | CAGTGGGGCGGTTGTTAT | 18 | 0.01 | 67.9603 | 12.73048 | 6.20E-06 | Up |
| miR-6313-y | TGCTGTGAAGTTTTGATT | 18 | 0.01 | 34.2258 | 11.74087 | 0.000177 | Up |
| miR-2965-y | AGGACTGCTACAGAGAGCA | 19 | 0.01 | 29.1699 | 11.51026 | 1.18E-06 | Up |
| miR-8462-x | ATTAATTTGATAAGTTATA | 19 | 0.01 | 18.28523 | 10.83646 | 4.28E-05 | Up |
| miR-7338-y | TTTAGCTGGTTTGTCAAGA | 19 | 0.01 | 9.559467 | 9.900786 | 2.31E-05 | Up |
| miR-3654-y | GCGACTGGAAAAGCTGAA | 18 | 0.01 | 6.910967 | 9.432744 | 2.64E-06 | Up |
| miR-3720-x | TACGGTGATGAGTTTAAA | 18 | 24.48687 | 63.4207 | 1.372946 | 0.023317 | Up |
| novel-m0019-5p | AGTCTCGATCGAGACATGTGA | 21 | 8.374933 | 0.01 | -9.70993 | 1.58E-06 | Down |
| novel-m0003-3p | TGGTGATATGTGTATATACTGATT | 24 | 8.917333 | 0.01 | -9.80047 | 8.91E-05 | Down |
| miR-92-x | TTGGGCGGGGTGTCCGTGC | 19 | 31.94557 | 0.01 | -11.6414 | 0.001613 | Down |
